# The Evolution of Host Specialization in an Insect Pathogen

**DOI:** 10.1101/2021.10.11.463986

**Authors:** Elisa Visher, Lawrence Uricchio, Lewis Bartlett, Nicole DeNamur, Aren Yarcan, Dina Alhassani, Mike Boots

## Abstract

Niche breadth coevolution between biotic partners underpins theories of diversity and co-existence and influences patterns of disease emergence and transmission in host-parasite systems. Despite these broad implications, we still do not fully understand how the breadth of parasites’ infectivity evolves, the nature of any associated costs, or the genetic basis of specialization. Here, we serially passage a granulosis virus on multiple inbred populations of its *Plodia interpunctella* host to explore the dynamics and outcomes of specialization. In particular, we collect time series of phenotypic and genetic data to explore the dynamics of host genotype specialization throughout the course of experimental evolution and examine two fitness components. We find that the *Plodia interpunctella* granulosis virus consistently evolves increases in overall specialization, but that our two fitness components evolve independently such that lines specialize in either productivity or infectivity. Furthermore, we find that specialization in our experiment is a highly polygenic trait best explained by a combination of evolutionary mechanisms including conditionally positive fitness asymmetries and mutation accumulation. These results are important for understanding the evolution of specialization in host-parasite interactions and its broader implications for co-existence, diversification, and infectious disease management.

## Introduction

The question of why some species are specialists and others are generalists has been central to evolutionary biology since its inception (Darwin, 1859). This co-existence of strategies is commonly explained by there being some cost to generalism such that specialists are favored under certain ecological conditions (Futuyma and Moreno, 1988) because “jacks-of-all-trades are the masters of none” (MacArthur 1984). The theory of costly generalism has been extensively applied in the host-parasite eco-evolutionary literature to explain parasite niche breadth and specialization at the levels of both host species and host genotype (Gandon and Poulin, 2004; Osnas and Dobson, 2012; Regoes et al., 2000). Niche breadth at the level of host species has important implications for pathogen emergence (Guth et al., 2019) and species co-existence (Connell, 1971; Janzen, 1970); while niche breadth and specialization at the genotype level underpins the monoculture effect (Elton, 1958), local adaptation (Kawecki and Ebert, 2004), and the Red Queen Hypothesis of Sex (Jaenike, 1978).

Despite these broad implications for niche breadth evolution in antagonistic coevolutionary systems, there is still debate about whether costs to niche breadth are, in fact, universal and what the dominant genetic mechanisms driving such costs would be (Jaenike, 1990; Remold, 2012). Several mechanisms for the evolution of specialization have been proposed. The classic trade-off hypothesis expects that increases in fitness on one host negatively trade-off with fitness on foreign hosts (Levins, 1968; Regoes et al., 2000). These strict negative trade-offs are not universal though, so several additional theories have been proposed, including host specialization due to weakly positive or neutral genetic correlations leading to asymmetrical fitness gains (Fry, 1996) and host specialization due to the accumulation of deleterious mutations on alternate hosts (Kawecki, 1994; Whitlock, 1996). The number of genes involved in specialization could also vary so that it is driven by few mutations of large effect or by many mutations of small effect.

Experimental evolution approaches have been used to explore host genotype specialization with great success, but in a relatively limited number of host-parasite systems including mice and RNA virus (Kubinak et al., 2012), mosquitos and microsporidia (Legros and Koella, 2010), daphnia and bacteria (Little et al., 2006), protists and bacteria (Nidelet and Kaltz, 2007), *C. elegans* and bacteria (Schulte et al., 2011), and wheat and fungus (Zhan et al., 2002). Generally, these studies find that serial passage on a single host genotype increases fitness on that host genotype while decreasing or at least resulting in smaller fitness gains on other genotypes.

However, there has been limited empirical exploration of the genetic mechanisms of such specialism. Specifically, these experiments have not explored the number of potential mutations involved in host genotype specialization or the genetic mechanisms of such specialization. Remold et al. (2008) do directly test of the genetic mechanism of specialization in a host-pathogen system to find a mix of antagonistic pleiotropy and mutation accumulation depending on evolutionary history, but their experiment looks at adaptation to different host cell lines rather than host genotype. A better understanding of the genetics of specialization is important because the potential number of mutational solutions has implications for population genetics theory on the speed of evolution and on how epistasis can promote costly generalism (Remold, 2012). Furthermore, different genetic mechanisms driving costly generalism may create divergent predictions for eco-evolutionary theory (Visher and Boots, 2020).

In this paper, we explore the evolutionary dynamics of host genotype specialization in the *Plodia interpunctella* (Hübner) and Plodia interpunctella granulosis virus (PiGV) laboratory model system. *Plodia interpunctella*, the Indian meal moth, is a stored grain pest that has been extensively used to characterize trade-offs and test eco-evolutionary dynamics in the lab (Bartlett et al., 2020, 2018; Boots, 2011; Boots and Mealor, 2007). We experimentally evolve virus populations to determine whether PiGV evolves to specialize on familiar host genotypes, collect multiple fitness metrics at multiple time points to explore the phenotypic dynamics of specialization, and sequence virus populations at multiple time points to explore the genetic mechanisms of specialization. We find that serially passaging virus leads to consistent increases in specialization on familiar host genotypes through the course of experimental evolution, and that specialization can occur in multiple fitness components. MCMC-based inference analysis of time series data shows that this specialization is not driven by few mutations of large effect (Schraiber et al., 2016). Combining these lines of evidence suggests that a combination of genetic mechanisms is likely to explain specialization in our system.

## Methods

### Study System

Our study system is *Plodia interpunctella* (Hübner), the Indian meal moth, and the Plodia interpunctella granulosis virus (PiGV). *Plodia interpunctella* is a pest that lives in grain stores (Mohandass et al., 2007). During its five larval instar stages, it develops within its food medium before pupating and emerging into an adult moth. For this experiment, we use inbred lines previously generated in Bartlett et al. (2018). These lines were made by mating individual brother-sister pairs for more than 27 generations. At this point, inbred populations should represent near-clonal populations of a single genotype that was randomly selected from the genetically diverse founder population via drift.

Plodia interpunctella granulosis virus (PiGV) is a dsDNA baculovirus that is an obligate killer (Vail and Tebbets, 1990). The natural life cycle is as follows: a larvae ingests virions in the occlusion body form, the virions shed their protein coats and infect gut epithelial cells, the virions either pass through the gut to establish a successful infection or are cleared during molting (freeing the larvae to carry out the rest of their life history), the virus begins to proliferate through the entire body of the larvae, and, once at a critical mass, packages into the protein-coated occlusion body form and kills its host (Rohrmann, 2013). It can then be transmitted to susceptible larvae when they cannibalize infected cadavers and ingest occluded virus. Critically, the virus must kill its host in order to transmit and larvae can only pupate and become adult moths if they were not successfully infected (Boots and Begon, 1993).

### Host Selection and Maintenance

We selected three inbred *Plodia interpunctella* populations with similar overall levels of resistance for this experiment, as measured by a preliminary resistance assay of all twelve of the inbred populations (unpublished data). The chosen inbred populations (Lines 2, 9, and 17) represent genotypes with similar medium overall levels of resistance (Supplemental Table 1). Populations of these genotypes were maintained in the absence of the virus as in (Bartlett et al., 2020) (See Supplemental Methods for details).

### Setting Up Experimental Evolution

Virus evolution was initiated with a single genetically diverse virus stock that we diluted to a passaging dose that would cause high mortality (~7.5×10^8 occlusion bodies per mL). We counted the concentration of this passaging dose on a Petroff-Hauser counting chamber with a darkfield microscope at 400x magnification. This dilution was combined with 2% sucrose (ThermoFisher Scientific, U.S.A.) and 0.2% Coomassie Brilliant Blue R-250 dye (ThermoFisher Scientific, U.S.A.). The sucrose encourages the larvae to consume the virus solution and the dye allows us to recognize larvae that have consumed half their body length of virus solution and are therefore considered successfully inoculated.

We set up three replicate evolving lines of virus on each of the three inbred host genotypes (See Supplemental Figure 1 for passaging scheme). For each virus line, we collected 100 third instar larvae of the appropriate genotype in a petri dish and starved them under a damp paper towel for 2 hours. We then syringed tiny droplets of our virus-sucrose-dye solution onto the petri dish for the larvae to orally ingest. After about an hour, we moved 50 successfully inoculated larvae into two 25-cell compartmentalized square petri dishes (ThermoFisher Scientific, U.S.A.) with standard food. The grid plates were then transferred to a single incubator for 20 days.

### Serial Passage

After 20 days, we harvested virus from each virus line under sterile conditions by collecting up to 10 virus killed cadavers per line and transferring these to sterile 15 mL disposable tissue grinders (ThermoFisher Scientific, U.S.A.). Infected larvae were recognizable by their opaque, chalky, white coloration. We were not able to collect 10 infected cadavers from all virus lines at all passages, so, when we could not find 10 infected cadavers, we collected every infected cadaver that we could find (See Supplemental Table 2). To extract virus from infected cadavers, we added 2mL of sterile DI water to the tissue grinders and homogenized the solution until all cadavers had been thoroughly crushed. We then transferred 1mL of the supernatant to a sterile 1.5mL Eppendorf tube and centrifuged the solution for 1 minute at 3,000 rpm to remove larger particulate matter from the supernatant. We transferred 600uL of this solution to a sterile 1.5 mL Eppendorf and centrifuged this for 3 minutes at 13,000 rpm to pellet the virus. We removed the supernatant from the pellet and resuspended in 1mL sterile water.

After extracting the virus, we diluted the solution 10x and added 600uL of the dilution to a .65 micron filter spin column (Millipore Sigma, U.S.A.) that we centrifuged at 13,000rpm for 3 minutes to semi-purify the virus of possible bacterial and fungal contaminants (for method details see Supplemental Table 3). We counted each of the semi-purified virus solutions as above and diluted them to the passaging dose concentration of ~7.5×10^8 occlusion bodies per mL in 2% sucrose and .2% dye to form our final passaging solutions for each virus line. These virus dilutions were then used to infect the next set of third instar larvae of the appropriate genotype following the protocol above. Virus was serially passaged for nine passages.

### Assaying

We assayed each virus line at multiple passages to track evolution over the course of the experiment. We assayed the starting population of virus as well as virus harvested from passages 1, 4, 6, and the final passage 9. By assaying passage 1, we were able to check for the ecological effects of strain sorting from our genetically diverse starting virus population. For each assay, we inoculated all 3 host genotypes with all 9 virus lines at both the passaging dose and 10% of the passaging dose. We inoculated 25 larvae for each host genotype × virus line × dose combination using the standard inoculation protocol above. Because of time constraints, inoculations for each passage were conducted across three days with one host genotype each day being inoculated with all of the virus lines. By assaying all the virus populations from each of the evolutionary histories on all of the host genotypes, we were able to measure how the evolving virus line changed in fitness on the familiar (the genotype that the virus evolved on) and foreign (genotypes that the virus was unexposed to) host genotypes.

After 20 days, we froze the grid plates and counted the number of infected and uninfected individuals in each grid. We collected all the infected larvae from each assay grid that had been inoculated with the higher dose and froze them in a pooled sample per grid plate. We extracted virus from these samples via tissue grinding and the two centrifugation steps (without filtering) and counted the virus in a Petroff-Hauser counting chamber as above. From these virus counts and the number of infected larvae, we were able to determine how many occlusion bodies each virus line produced per infected cadaver on average when infecting each host genotype.

Finally, we multiplied the average number of virions produced per infected cadaver by the proportion of larvae infected to get a composite measure of fitness for each virus line on each host genotype.

### Sequencing and Variant Calling

The ancestor virus population and virus populations for each line at the four assayed time points (37 samples) were next prepared for sequencing. First, extracted occlusion bodies (OBs) were rinsed in 0.1% SDS and purified in a Percoll gradient as in (Gilbert et al., 2014). OBs were then dissolved in 0.5M Na_2_CO_3_ and DNA was extracted with a QIAamp DNA kit. Library preparation and sequencing was conducted at the UC Berkeley QB3 center on non-amplified DNA. 150bp paired end libraries were generated with Kapa Biosystems library preparation kits and multiplexed to run on one lane of an Illumina MiSeq platform. Reads were then de-multiplexed and aligned to the PiGV reference genome [GenBank: KX151395] using bowtie2 (Harrison et al., 2016; Langmead and Salzberg, 2012). The resulting alignments for each sample had 99.99-100% genome coverage, 51-100 mean coverage depth, and 40.2-41 mean MapQ scores. Variant calls were made using VarScan (Koboldt et al., 2012) in Galaxy, calling indels and SNPs separately using a minimum coverage of 20 and a minimum alternate allele read count of 2. The Galaxy history can be viewed here: https://usegalaxy.org/u/evisher/h/specialization-sequence-variants.

### Phenotypic Assay Data Analyses

We analyzed all phenotypic assay data using a linear mixed modelling approach in R (v.3.4.4 – “Someone to Lean On”) (R Core Team 2018), accounting for our mixed design using the ‘afex’ package (Singmann et al., 2019) which relies on the ‘lme4’ linear mixed modelling engine (Bates et al. 2015). Where appropriate, we followed this with post-hoc testing using the ‘emmeans’ package (Lenth, 2019) to identify pairwise differences between levels of interacting fixed effects, with *p*-values corrected for multiple comparisons using a Bonferroni correction. Our response variables were either fitness, infectivity, or productivity of the virus line. We used a binomial error structure for models of infectivity and poisson error structures for models of productivity and fitness.

The first part of our analysis looked at data from the end of the evolution experiment (passage 9). We tested for an effect of specialization by using a ‘self’ factor that was either true (virus was assayed on same host genotype it was evolved on) or false (virus was assayed on a host genotype it was not evolved on). We first included this as a fixed effect alongside ‘assay genotype’ and ‘evolution genotype’ (the host genotype used for the assay and that the virus was evolved on, respectively). In the case of the ‘infectivity’ data analysis, ‘dose’ was also included as a fixed effect. Our random effects were ‘evolution genotype’ and ‘virus line’, with ‘virus line’ nested under ‘evolution genotype’ to account for our experimental structure. We further used the same modelling approach to test for a correlation between a virus line’s virion production and its infectivity by including virion production as an additional fixed predictor in a separate model of infection likelihood at the highest dose.

We further analyzed the effect of ‘self’ by including it as a fixed term in models the same as specified above, however with ‘virus line’ replacing ‘evolution genotype’ as a fixed effect (no change to the random effects), and a potential interaction between ‘self’ and ‘virus line’ to see if there were differences in the ability of each virus selection line to evolve any specialism. We used pairwise comparisons between lines to investigate these differences; further, we used these pairwise differences to test for a correlation between each virus line’s fitness on a familiar genotype and its fitness on a foreign genotype.

We also analyzed our fitness data across the whole experiment, including data for passages 0, 1, 4, 6, and 9, to interrogate how specialization evolved with time. We used the same general approach as detailed above, where fixed effects were ‘assay genotype’, ‘self’, ‘passage number’, and an interaction between ‘passage number’ and ‘self’. Our error structure included ‘evolution genotype’, ‘virus line’ and ‘passage number’, with ‘virus line’ nested under ‘evolution genotype’ as above, and ‘passage number’ nested under ‘virus line’ to account for multiple generations acting as repeated measures.

We used the ‘ggplot2’ (Wickham, 2009) package to plot graphs of our results. See supplement for annotated code.

### Variant Analysis

Variant frequencies were analyzed to: 1) identify genetic regions of variation in our population, 2) determine whether variant community composition was predicted by treatment, and 3) identify signatures of positive selection across the time series.

To determine regions of variation, we plotted variant frequencies against genome position, identified genome regions with high genetic variation, and compared these genetic regions to the annotated PiGV reference genome by hand to identify potentially interesting nearby genes (Harrison et al., 2016). To determine whether passage 9 variant community composition was predicted by treatment, we used constrained ordination analysis on Hellinger pre-transformed SNP frequencies using the ‘vegan’ package and performed a Monte Carlo permutation test to determine whether treatment significantly predicted SNP frequency variance amongst the virus populations (Oksanen et al., 2020). See supplement for annotated code.

Finally, to identify signatures of positive selection, we used an MCMC-based inference procedure to infer the strength of selection acting at variable positions in our genomic time series data (Schraiber et al., 2016). This software estimates selection coefficients given an observed frequency trajectory, accounting for uncertainty in true allele frequencies due to binomial sampling. While we knew the average virus population size within a single individual at the end of infection, we did not know the exact number of virus particles that founded each infection. Thus, we chose several possible demographic models based upon a range of inoculums (from 50 to 500 viral particles) and a range of growth rates (including “slow” and “fast” exponential processes with ~1.2-5 fold growth per generation) and calculated the harmonic mean of population size and the number of generations needed to reach 10^10^ particles for each scenario (Harpak and Sella, 2014). We then repeated our estimates of selection strength using each of these effective population sizes, which ranged from small to moderate (N_e_ = 425 to N_e_ = 6778). We call ‘significant’ alleles using the most conservative demographic model (N_e_=425) by a loose threshold, where the 90% HPD interval did not overlap 0. For details, see supplementary methods and annotated code.

## Results

### Specialization of Viruses at the Final Passage

After nine passages of experimental evolution, we find good evidence that viruses evolved to specialize on their familiar host genotype, indicated by a significantly positive effect of ‘self’ on viral infectivity (p = 0.01), productivity (p < 0.001), and overall fitness (p < 0.001) (See Figure 1). Therefore, the evolved virus lines infected relatively higher proportions of individuals, produced more virions per infection, and therefore had higher fitness when infecting the host genotype that they had evolved on than when infecting foreign host genotypes. As expected, we found a significant effect of ‘assay genotype’ (host) across all three response variables (p < 0.002 in all cases) and a significant positive effect of ‘dose’ (p < 0.001) when analyzing infection likelihood. We did not find a significant effect of ‘evolution genotype’ on infection likelihood (p = 0.69) or fitness (p = 0.12), meaning that specific host genotypes did not lead to the evolution of generally more infectious or higher fitness virus populations when averaged across all three assay genotypes. However, we did find a significant effect of ‘evolution genotype’ on viral productivity (p = 0.03), driven by the higher productivity of virus lines evolved on host genotype 17 (See Supplemental Model Tables).

**Figure 1.**
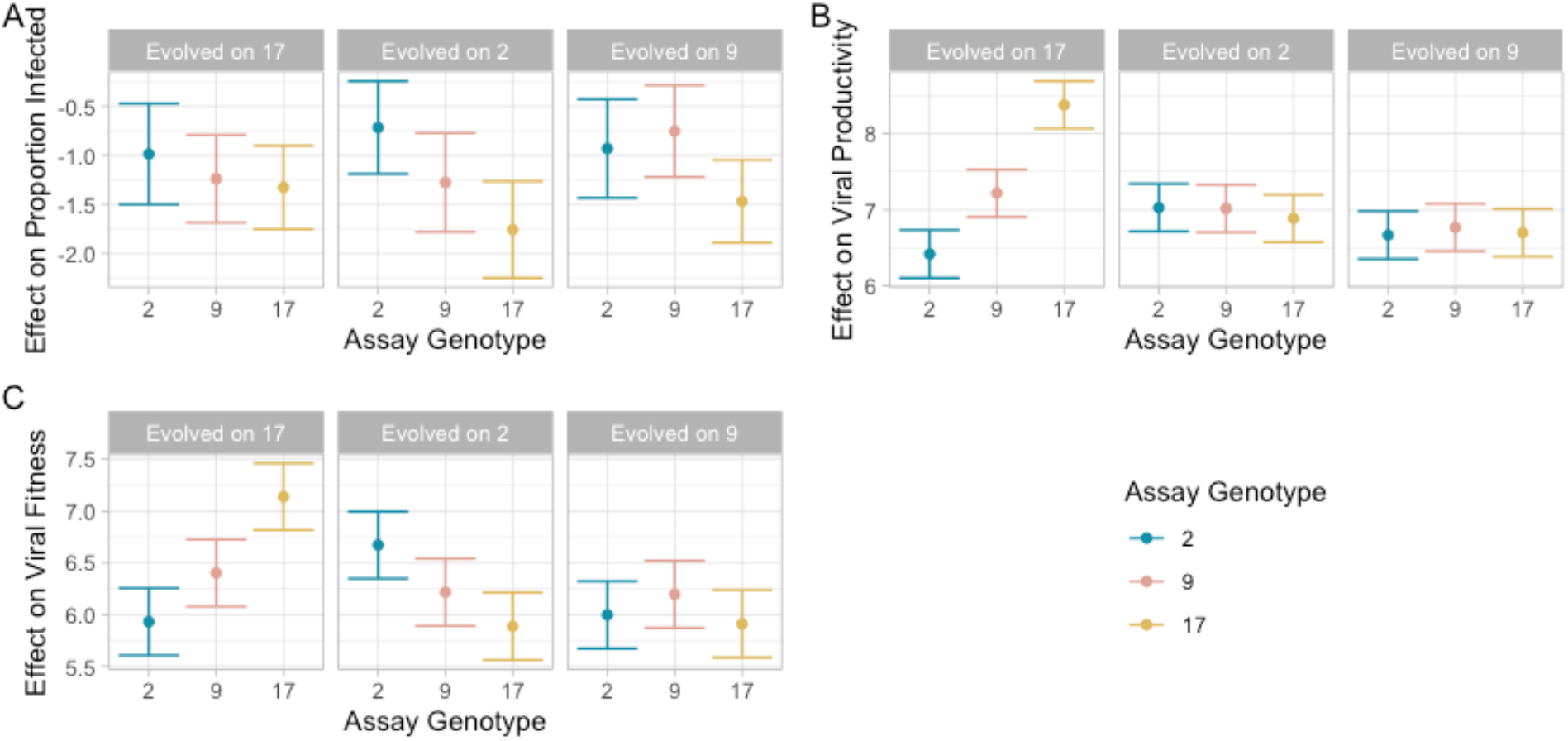
Specialization of virus at the end of the experiment. Paneled plots show the effect of the virus’s evolutionary history on its (a) infectivity, (b) productivity, and (c) composite fitness when infecting each of the assay lines. Fitness is the proportion infected x the average number of virions produced per infected cadaver. Virus lines are significantly more infective, productive, and fit on familiar genotypes. Y-axis effect sizes and errors are taken from the GLMM models using the ‘emmeans’ package.

We found evidence that virus lines differed both in overall fitness and in specialization when we tested for an effect of ‘virus line’ and an interaction between ‘virus line’ and ‘self’. We found a significant effect of ‘virus line’ and a significant interaction between ‘virus line’ and ‘self’ when analyzing both the fitness and viral productivity data (p < 0.001 in both cases), but not when analyzing the infectivity data (p = 0.06 for ‘virus line’, p = 0.25 for ‘virus line’:‘self’ interaction). That is, virus lines differed in their overall fitness and productivity (but not infectivity) across all the assay genotypes and, while lines generally showed higher fitness and productivity on their familiar host genotype, this effect varied significantly amongst lines.

Using a Bonferroni-corrected pairwise comparison analysis, we find that most lines differ from each other in fitness: only 4/36 pairwise comparisons between lines showed no significant differences in fitness on a familiar host, and only 8/36 showed no difference on a foreign host. The large majority of lines therefore differed from all other lines in their fitness on both hypothetical average familiar and foreign hosts. Only one pair of lines (9.2 and 2.2) showed no difference from one another on both foreign and familiar host genotypes. Looking at the effect of ‘self’ for each line, our pairwise comparisons also explain that the effect of ‘self’ differs between lines. Line 9.1 was significantly less fit on a familiar host compared to a foreign one (p < 0.001), while lines 9.2 and 2.2 showed no significant differences in their fitnesses on familiar compared to foreign hosts (p = 0.50 and p = 0.28 respectively). All other lines were significantly more fit on a familiar host than a foreign one, though there were differences in the magnitude of this effect. Notably, these virus line differences and their interaction with ‘self’ were due to increased virion production; lines didn’t significantly differ in their ability to infect an ‘average’ host and didn’t differ in how much more likely they were to infect a familiar host.

### Evolution of Specialization over Time

Our analysis of fitness data across all passages (0,1,4,6,9) showed a significant effect of passage number on virus fitness (p < 0.001), and a significant interaction between passage number and ‘self’ (p < 0.001). Virus lines decreased in fitness from passage 0 to passage 1 and generally increase in their overall fitness from passage 1 to passage 4, with no further meaningful overall change from passages 4-6 and 6-9. The initial decrease in fitness from passage 0 to passage 1 could be due to the different storage and extraction methods of the starting virus stock from experimental virus solutions, and analyzing fitness without passage 0 data confirms the passage 1-9 trends. However, viruses increase their fitness on their familiar host genotype **relative to their foreign host genotype** in every case (from 1-4, 4-6, and 6-9), suggesting either a gain in fitness on the familiar genotype, a loss of fitness on the foreign genotype, or both (see Figure 2). Interestingly, when we analyze the infectivity and productivity data separately, we do not see this steadily increasing effect of self across the time course of the experiment.

**Figure 2.**
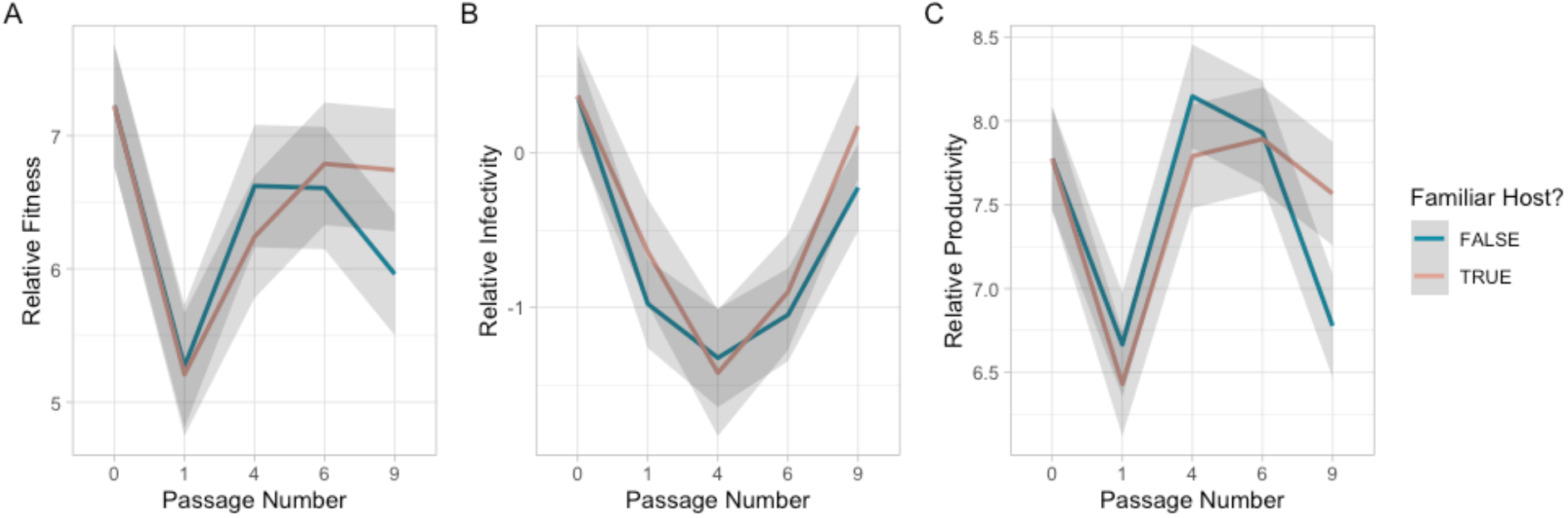
Evolution of specialization over time. Panelled plot showing the effect of whether the virus was assayed on its familiar host genotype (red) or on a foreign one (blue) on viral (a) infectivity, (b) productivity, and (c) fitness over time. Virus lines evolve significantly higher relative infectivity, productivity, and fitness on familiar lines over the experiment. Y-axis effect sizes and errors are taken from the GLMM models using the ‘emmeans’ package.

### Relationship between Virus Productivity and Infectivity

We find a significant negative correlation between virus productivity and infection likelihood (p < .001) in the passage 9 dataset. However, when we analyzed the full dataset with all passages, we find that virus productivity and infectivity were significantly positively correlated (p<.001), such that more productive virus lines are also more infective. Therefore, we fit and test a model with an interaction effect between ‘self’, ‘productivity’, and ‘passage number’ and find a significant interaction between these three metrics (p<.001) such that the direction of the relationship between viral productivity and infectivity changes from positive to negative depending on the passage number and whether the virus is infecting a familiar or foreign host (see Figure 3).

**Figure 3.**
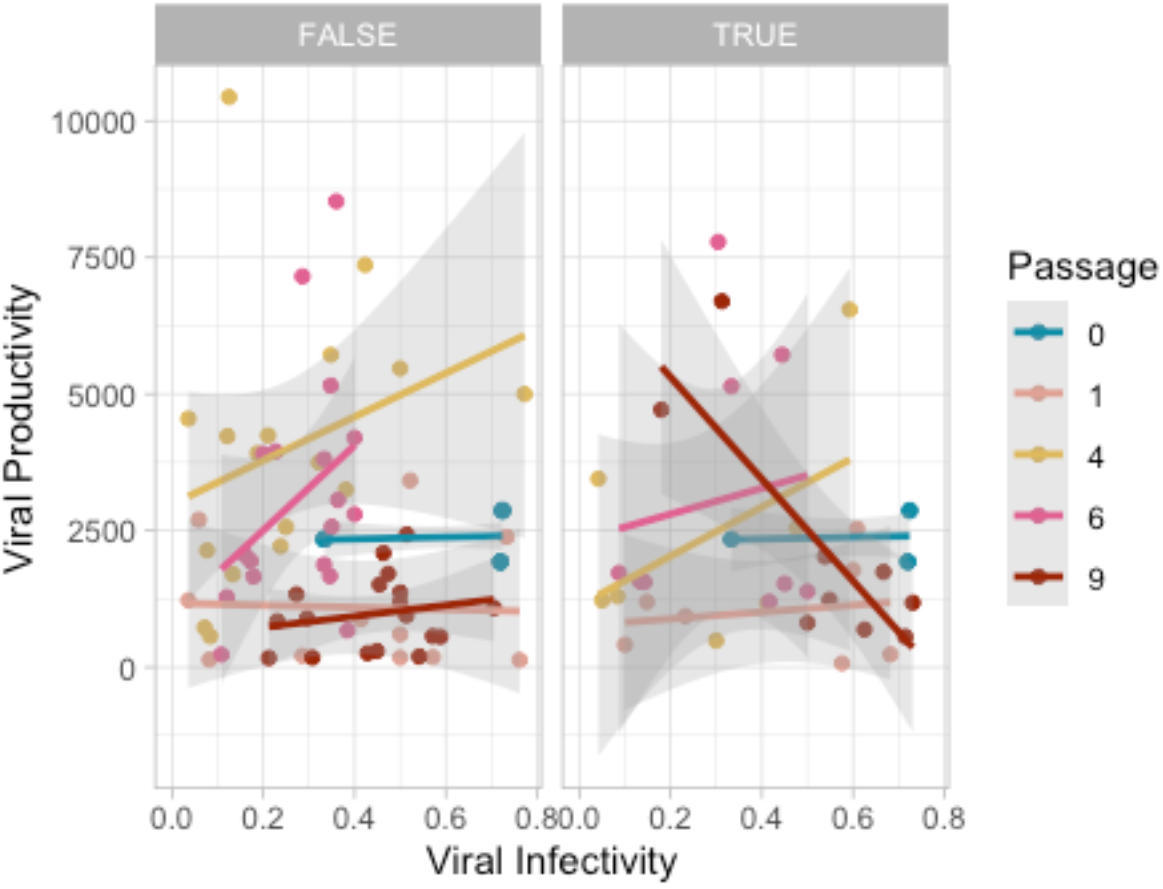
Relationship between Virus productivity and infectivity. Paneled plot showing the relationship between viral productivity and infectivity on both familiar (right panel) and foreign hosts (left panel) at each passage. Infectivity is defined as the proportion of hosts infected and productivity is the average number of virions produced per infected host. The direction of the relationship between viral productivity and infectivity significantly changes from positive to negative depending on the passage number and whether the virus is infecting a familiar or foreign host.

### Correlation between Fitness on Familiar and Foreign Hosts

We next determined the correlation between a virus line’s fitness on their familiar host genotype and on the foreign host genotypes. A negative correlation would mean that the virus lines with the highest fitness on their familiar genotype had the lowest fitness on foreign genotypes and indicate a strict trade-off. At the final passage of experimental evolution, there was no significant effect of fitness on familiar genotypes on fitness on foreign genotypes (t_7_ = − 0.48, p = 0.65). Extending this analysis to include passages 4, 6, and 9, we continue to find no significant correlation between fitness on familiar and foreign genotypes (t_23_ = 1.7, p = 0.11) (See Figure 4).

**Figure 4.**
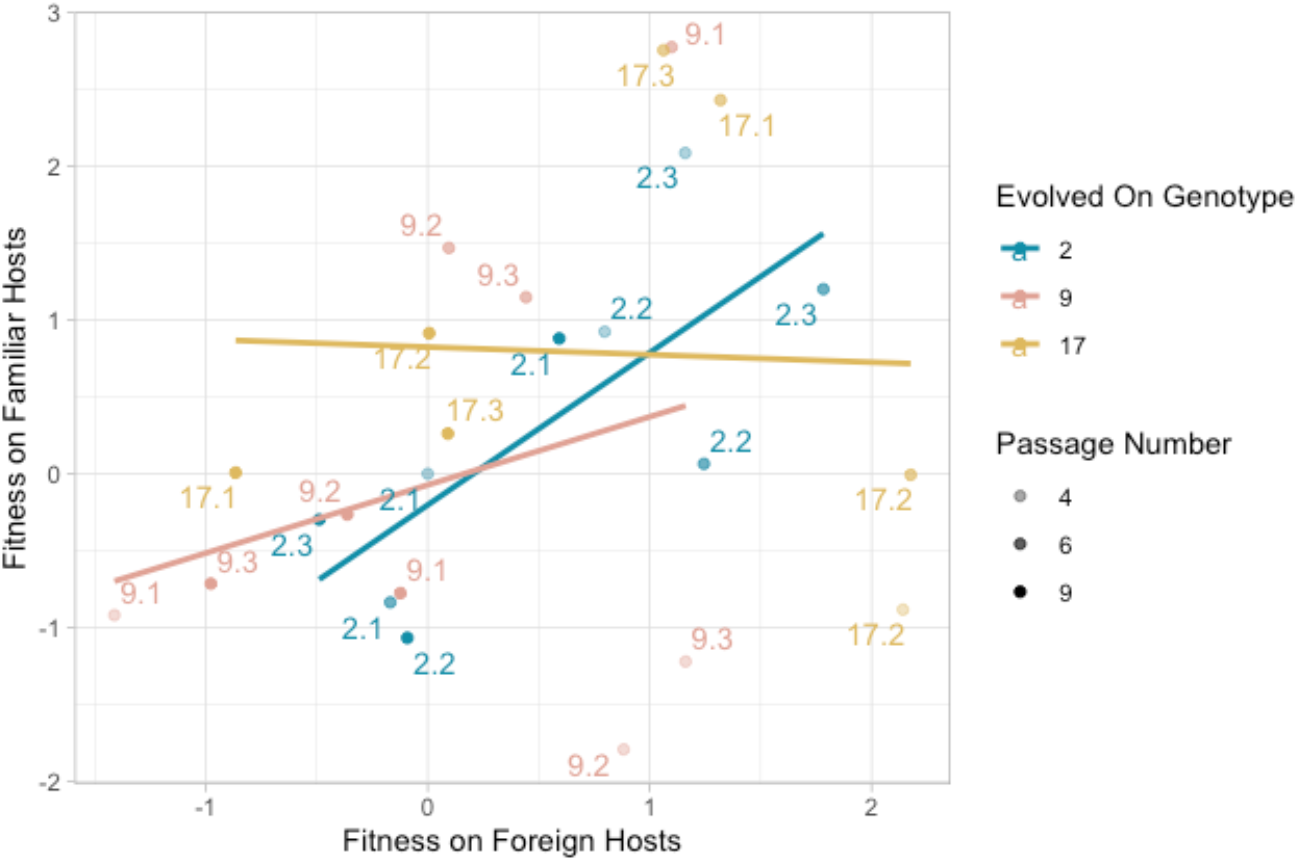
Correlation Between Fitness on Familiar and Foreign Hosts. Plot showing the correlation between each virus line’s fitness on familiar and foreign hosts at passages 4, 6, and 9. There are no significant trends between fitness on foreign and familiar hosts across the experiment. Effect sizes are taken from the GLMM models using the ‘emmeans’ package.

### Genetic Variation

Most variants are at low (<10%) frequencies, but there are several genomic regions that consistently have high genetic variation (Supplemental Figure 8). These regions correspond with several ORFs homologous with genes in AcMNPV that have known functions including occluded virus production, oral infection, time to kill, and host range (Supplemental Table 4) (Harrison et al., 2016; Rohrmann, 2019). We do not find that treatment significantly predicts variance in SNP community composition at passage 9 in constrained ordination analyses with permutation tests (25% variance explained, p=.15), indicating that treatment is not significantly predicting the frequencies of genetic variants.

Among the 114 alleles that were called as significant in the analysis of the *N_e_* = 540 model (Figure 5), we found a small subset that were called as significant in 2 or more biological replicates from the same treatment (Supplemental Table 5). A majority of these variants were indels (13 out of 20). Only two indel variants had selection signals that were specific to a particular condition (variants at positions 27789 and 20105, both of which were detected in multiple replicates of line 17). Other putatively selected variants were shared across virus populations from two or more of the inbred lines, suggesting they may represent adaptation to experimental conditions rather than adaptation to specific host genotypes. Among the 7 selection signals at SNPs, 3 were unique to specific host genotypes, including positions 27885, 33317, and 33358, which were detected in viruses evolved on lines 9, 17, and 17, respectively. In general, we note that the inferred selection coefficients are mostly indicative of weak positive selection. If we suppose an effective population size of 540, then the inferred values of *2Ns* indicate per-allele effects ranging from 0.0065 to 0.042.

**Figure 5.**
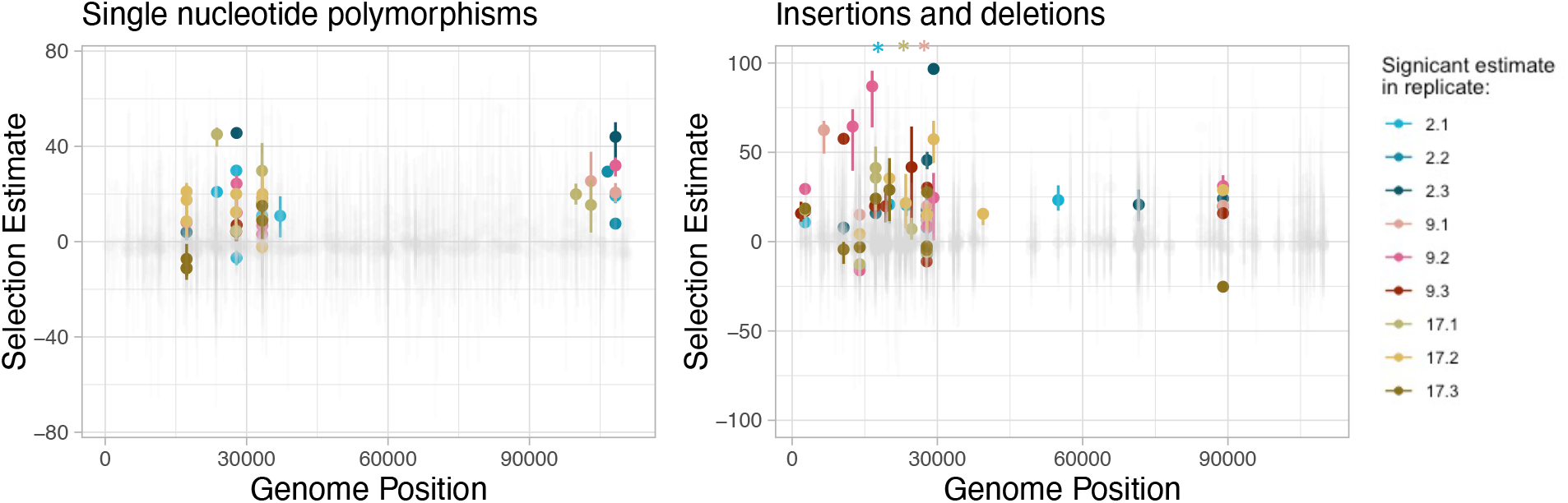
Genome-wide selection inferences for SNP and indel variants in each replicate. Significant variants (positive and negative) are colored by the replicate that they are detected in, and non-significant variants are light gray. Stars indicate selection estimates outside of the graph area. See supplemental figures for non-condensed versions of this figure.

## Discussion

Specialization is critical to many of our theories of coevolution and the maintenance of diversity (Futuyma and Moreno, 1988). In particular, specialization between parasites and their hosts is crucial for understanding patterns of disease emergence and spread (Woolhouse and Gowtage-Sequeria, 2005). Here, we use experimental evolution techniques to test whether a granulosis virus can evolve to specialize on specific genotypes of its moth host. We find that the virus evolved to specialize in infectivity, productivity, and overall fitness on familiar host genotypes, though the degree of specialization depends on the virus line.

A unique feature of our experiment is that we collect time series phenotypic and genetic data that allow us to explore the dynamics of specialization in novel ways. First, a key finding of our experiment is that the virus specializes on familiar host genotypes in both viral infectivity and productivity (Figure 1). Several previous similar studies have also measured multiple fitness components related to specialization to find that pathogens could variably specialize on parasite virulence and/or transmission (Kubinak et al., 2012; Legros and Koella, 2010; Nidelet and Kaltz, 2007; Zhan et al., 2002). However, none of these previous studies have examined the correlations between their fitness components across time.

With our phenotypic time series data, we can see that the relationship between our two fitness components, viral infectivity and productivity, is positive at the start of the experiment but, by passage 9, evolves be negative when infecting familiar hosts (Figure 3). This suggests that different virus lines are primarily being selected to increase specialization on familiar host genotypes in either viral productivity or viral infectivity. This finding highlights the importance of measuring multiple fitness components when pathogen populations can use many strategies to increase their overall fitness and suggests that researchers should be hesitant to conclude that specialization (or local adaptation) did not evolve.

Furthermore, our time series data allow us to test hypotheses about the dynamics of specialization within our system. At passages 1 and 4, virus lines are slightly less fit on their familiar host genotype, suggesting that selection in our experiment was not dominated by the ecological sorting of virus genotypes from the genetically diverse starting population. Instead, the steadily increasing effect of self across the course of the experiment suggests that evolutionary processes were more important.

Next, we can ask questions about the number of potential genes involved in specialism evolution. If specialization were to be driven by few mutations of large effect, we would expect to see some degree of genetic parallelism in replicates and strong signatures of positive selection (if specialization is not driven by mutation accumulation). If there were many genetic options for specialization, we would less expect to see the phenotypic parallelism of the experiment reflected at the genetic level and selection on any one variant would be weaker. We also might expect to see slower changes in phenotypic time series data.

In our experiment, the evidence indicates that specialization was driven by many variants of small effect. Our phenotypic time series data for individual virus lines suggests that specialization slowly increases and wanders, pointing against the fixation of large effect mutations (Supplemental Figure 2). This is indeed confirmed by our sequencing data in which we did not observe any clear signals of selective sweeps where low frequency alleles swept to high frequency. Given the relatively high depth of coverage of our samples and the quality of the sequencing data, it is unlikely that we failed to detect many (if any) sweeps. Furthermore, our selection analysis does not identify any variants with strong parallel signatures of selection across replicates.

Second, we can ask questions about whether specialization is driven by antagonistic pleiotropy, conditionally positive adaptation resulting in fitness asymmetries, or mutation accumulation in alternate environments. If specialization were to be driven by antagonistic pleiotropy, we would expect to see that the most fit replicates on the familiar host are the least fit on the foreign host and that positive selection acts on variants. We would not have clear predictions for how total fitness across all the genotypes would change over time as this would depend on the symmetry of the trade-off shape. In the case of conditionally positive alleles resulting in fitness asymmetries between familiar and foreign hosts, we would expect to see slightly positive or neutral fitness correlations between familiar and foreign hosts, positive selection on variants, and overall increases in total fitness across all the genotypes. In the case of mutation accumulation, we would expect negative fitness correlations between familiar and foreign hosts (the most specialized are those that are worst on foreign hosts), no evidence of positive selection since MA is driven by drift, and overall decreases in total fitness across all the genotypes. Of course, these mechanisms are not exclusionary, especially in our case where many variants can affect specialization. These predictions may therefore be muddied if multiple mechanisms are driving specialization. Additionally, any directional fitness changes to overall experimental conditions might hamper our ability to fully assess whether fitness correlations between genotypes are positive or negative (as some replicates may just be the ‘most adapted’ to the general environment) and our ability to assess changes in total fitness across genotypes in the system.

We did not find a significant correlation between fitness on familiar and foreign hosts, hampering our abilities to make conclusions about the mechanisms driving specialization. However, we can observe from Figure 4 that the sign of the correlation between fitness on familiar and foreign hosts may differ depending on the evolutionary background of the virus line. There appears to be a neutral or slightly negative correlation between fitness on host genotype 17 and foreign genotypes, suggesting that specialization on this host could be consistent with any mechanism. However, correlations between fitness on familiar and foreign host genotypes are positive for virus specializing on host genotypes 2 and 9. This suggests specialization driven by asymmetric conditional positivity. Nonetheless, the possibly neutral or negative correlations between fitness on host genotype 17 and foreign genotypes and the possibly positive correlations between fitnesses on host genotypes 2 and 9 and foreign genotypes would suggest that it is likely that multiple mechanisms contribute to specialization in our system.

From our sequence analysis, we do not see evidence of strong, parallel positive selection on any variants. We observe many instances of subtle frequency differentiation during the course of the experiment, which seems a likely candidate to explain the genetic mechanism for adaptation. This subtle frequency differentiation could be due to polygenic adaptation (in which frequency changes at many alleles of weak effect on a selected trait can have a substantial effect in concert) or selective interference (in which one clone fails to fix because a higher fitness clone invades and outcompetes it before fixation occurs). If the polygenic effects of these positively selected alleles are enough to explain the phenotypic specialization, then this would point towards specialization driven by weak positive selection on many alleles with pleiotropic or conditionally positive effects. However, if they are not, specialization could still be driven by mutation accumulation in alternate environments as this is a drift, not selection, based process.

Finally, the total fitness of virus lineages across all host genotypes does not increase continuously though the experiment. Total fitness does increases from passage 1 to passage 4 but plateaus from passage 4 to passage 9, which is also when we see our largest changes in specialization. This would suggest that antagonistic pleiotropy or a balance of conditional positivity and mutation accumulation is driving specialization. It also suggests that PiGV quickly reached a point of being fairly well adapted to experimental conditions so that directional selection to overall experimental conditions is less likely to obscure patterns resulting from specialization. However, a caveat to these trends in total fitness is that our assay scheme was designed to best test the changes in relative fitness on different genotypes over time and so assayed viruses from different passages on different days. Therefore, these trends in total fitness (but not relative fitness) might be confounded by random day effects.

In this experiment, we have shown that Plodia interpunctella granulosis virus can evolve to specialize on specific genotypes of its host and that specialization is not driven by strong selection on few alleles. However, we cannot precisely determine the evolutionary mechanism of this specialization. Putting our evidence together, it seems most likely that the evolution of specialization in our experiment is driven by many genetic variants and by multiple mechanisms. Most parsimoniously, a combination of weakly positive fitness asymmetries and mutation accumulation in alternate environments could explain how we have positive fitness correlations across familiar and foreign hosts without having increases in overall fitness. If conditionally positive fitness asymmetries are driven by many variants of small effect, it would explain the weak signatures of selection and lack of genetic parallelism in our sequence analysis. Then, these positive asymmetries combined with the drift-driven accumulation of conditionally deleterious variants (mutation accumulation) could have collectively driven specialization while their opposing effects on total fitness would result in no overall fitness changes.

In conclusion, we used an experimental evolution approach to determine whether a baculovirus could evolve to specialize on specific genotypes of its moth host. We find that virus does evolve higher infectivity, productivity, and overall fitness on familiar host genotypes. This specialization may be driven by a combination of conditionally positive alleles leading to fitness asymmetries and mutational accumulation on foreign host genotypes. Time series data shows that specialization in overall fitness evolves directionally over the time course of the experiment and that the different fitness components of virus lineages may be independently selected on. Our results demonstrate that gene-by-gene interactions are evolvable in the *Plodia interpunctella* and PiGV model system and suggests that the system has promise for experiments on the ecological conditions that shape selection on specialization and niche breadth.

## Supporting information

Supplemental model tables

## Author Contributions

EV and MB designed the experiment. EV, ND, AY, and DA collected data. EV, LB, and LU analyzed the data. EV, LB, LU, and MB wrote this manuscript.

## Acknowledgements

We would like to thank Annika McBride, Yazmin Haro, and Edith Lai for assistance with laboratory work. We would like to thank Britt Koskella for help designing the experiment. We would like to thank Britt Koskella and Bree Rosenblum for comments on the manuscript. EV acknowledges support from an NSF GRFP DGE 1752814 and the ASN George Gilchrist Student Research Award, and MB acknowledges support from NIH/R01-GM122061-03 and BBSRC BB/L010879/.

## Data Accessibly

All data presented in this study will be made available at a suitable repository following manuscript publication, and /or made available in the supplementary materials, alongside an annotated R script used for analysis.

## Works Cited

Bartlett, L.J., Visher, E., Haro, Y., Roberts, K.E., Boots, M., 2020. The target of selection matters: An established resistance—development-time negative genetic trade-off is not found when selecting on development time. J. Evol. Biol. 33, 1109–1119. https://doi.org/10.1111/jeb.13639

Bartlett, L.J., Wilfert, L., Boots, M., 2018. A genotypic trade-off between constitutive resistance to viral infection and host growth rate. Evolution 72, 2749–2757. https://doi.org/10.1111/evo.13623

Bates, D., Maechler, M., Bolker, B., Walker, S., 2015. Fitting Linear Mixed-Effects Models using lme4. J. Stat. Softw. 67, 1–48. https://doi.org/10.18637/jss.v067.i01.

Boots, M., 2011. The Evolution of Resistance to a Parasite Is Determined by Resources. Am. Nat. 178, 214–220. https://doi.org/10.1086/660833

Boots, M., Begon, M., 1993. Trade-Offs with Resistance to a Granulosis Virus in the Indian Meal Moth, Examined by a Laboratory Evolution Experiment. Funct. Ecol. 7, 528–534. https://doi.org/10.2307/2390128

Boots, M., Mealor, M., 2007. Local Interactions Select for Lower Pathogen Infectivity. Science 315, 1284–1286. https://doi.org/10.1126/science.1137126

Connell, J.H., 1971. On the role of natural enemies in preventing competitive exclusion in some marine animals and in rain forest trees. Dyn. Popul. 298, 312.

Darwin, C., 1859. On the origin of species, 1859. Routledge.

Elton, C.S., 1958. The ecology of invasions by animals and plants. Ecol. Invasions Anim. Plants.

Fry, J.D., 1996. The Evolution of Host Specialization: Are Trade-Offs Overrated? Am. Nat. 148, S84–S107.

Futuyma, D.J., Moreno, G., 1988. The Evolution of Ecological Specialization. Annu. Rev. Ecol. Syst. 19, 207–233. https://doi.org/10.1146/annurev.es.19.110188.001231

Gandon, S., Poulin, R., 2004. Evolution of multihost parasites. Evolution 58, 455–469. https://doi.org/10.1554/03-390

Gilbert, C., Chateigner, A., Ernenwein, L., Barbe, V., Bézier, A., Herniou, E.A., Cordaux, R., 2014. Population genomics supports baculoviruses as vectors of horizontal transfer of insect transposons. Nat. Commun. 5, 1–9. https://doi.org/10.1038/ncomms4348

Guth, S., Visher, E., Boots, M., Brook, C.E., 2019. Host phylogenetic distance drives trends in virus virulence and transmissibility across the animal–human interface. Philos. Trans. R. Soc. B Biol. Sci. 374, 20190296. https://doi.org/10.1098/rstb.2019.0296

Harpak, A., Sella, G., 2014. Neutral Null Models for Diversity in Serial Transfer Evolution Experiments. Evolution 68, 2727–2736. https://doi.org/10.1111/evo.12454

Harrison, R.L., Rowley, D.L., Funk, C.J., 2016. The Complete Genome Sequence of Plodia Interpunctella Granulovirus: Evidence for Horizontal Gene Transfer and Discovery of an Unusual Inhibitor-of-Apoptosis Gene. PLOS ONE 11, e0160389. https://doi.org/10.1371/journal.pone.0160389

Jaenike, J., 1990. Host Specialization in Phytophagous Insects. Annu. Rev. Ecol. Syst. 21, 243–273. https://doi.org/10.1146/annurev.es.21.110190.001331

Jaenike, J., 1978. A hypothesis to account for the maintenance of sex within populations. Evol Theory 3, 191–194.

Janzen, D.H., 1970. Herbivores and the Number of Tree Species in Tropical Forests. Am. Nat. 104, 501–528. https://doi.org/10.1086/282687

Kawecki, T.J., 1994. Accumulation of Deleterious Mutations and the Evolutionary Cost of Being a Generalist. Am. Nat. 144, 833–838.

Kawecki, T.J., Ebert, D., 2004. Conceptual issues in local adaptation. Ecol. Lett. 7, 1225–1241. https://doi.org/10.1111/j.1461-0248.2004.00684.x

Koboldt, D.C., Zhang, Q., Larson, D.E., Shen, D., McLellan, M.D., Lin, L., Miller, C.A., Mardis, E.R., Ding, L., Wilson, R.K., 2012. VarScan 2: Somatic mutation and copy number alteration discovery in cancer by exome sequencing. Genome Res. 22, 568–576. https://doi.org/10.1101/gr.129684.111

Kubinak, J.L., Ruff, J.S., Hyzer, C.W., Slev, P.R., Potts, W.K., 2012. Experimental viral evolution to specific host MHC genotypes reveals fitness and virulence trade-offs in alternative MHC types. Proc. Natl. Acad. Sci. 109, 3422–3427. https://doi.org/10.1073/pnas.1112633109

Langmead, B., Salzberg, S.L., 2012. Fast gapped-read alignment with Bowtie 2. Nat. Methods 9, 357–359. https://doi.org/10.1038/nmeth.1923

Legros, M., Koella, J.C., 2010. Experimental evolution of specialization by a microsporidian parasite. BMC Evol. Biol. 10, 159. https://doi.org/10.1186/1471-2148-10-159

Lenth, R., 2019. emmeans: Estimated Marginal Means, aka Least-Squares Means. R package version 1.3.5.1.

Little, T.J., Watt, K., Ebert, D., 2006. Parasite-Host Specificity: Experimental Studies on the Basis of Parasite Adaptation. Evolution 60, 31–38. https://doi.org/10.1111/j.0014-3820.2006.tb01079.x

MacArthur, R.H., 1984. Geographical Ecology: Patterns in the Distribution of Species. Princeton University Press.

Mohandass, S., Arthur, F.H., Zhu, K.Y., Throne, J.E., 2007. Biology and management of Plodia interpunctella (Lepidoptera: Pyralidae) in stored products. J. Stored Prod. Res. 43, 302–311. https://doi.org/10.1016/j.jspr.2006.08.002

Nidelet, T., Kaltz, O., 2007. Direct and Correlated Responses to Selection in a Host–Parasite System: Testing for the Emergence of Genotype Specificity. Evolution 61, 1803–1811. https://doi.org/10.1111/j.1558-5646.2007.00162.x

Oksanen, J., Blanchet, F.G., Friendly, M., Kindt, R., Legendre, P., McGlinn, D., Minchin, P.R., O’Hara, R.B., Simpson, G.L., Solymos, P., Stevens, M.H.H., Szoecs, E., Wagner, H., 2020. vegan: Community Ecology Package.

Osnas, E.E., Dobson, A.P., 2012. Evolution of Virulence in Heterogeneous Host Communities Under Multiple Trade-Offs. Evolution 66, 391–401. https://doi.org/10.1111/j.1558-5646.2011.01461.x

Regoes, R.R., Nowak, M.A., Bonhoeffer, S., 2000. Evolution of Virulence in a Heterogeneous Host Population. Evolution 54, 64–71. https://doi.org/10.1111/j.0014-3820.2000.tb00008.x

Remold, S.K., 2012. Understanding specialism when the jack of all trades can be the master of all. Proc. R. Soc. B Biol. Sci. 279, 4861–4869. https://doi.org/10.1098/rspb.2012.1990

Remold, S.K., Rambaut, A., Turner, P.E., 2008. Evolutionary Genomics of Host Adaptation in Vesicular Stomatitis Virus. Mol. Biol. Evol. 25, 1138–1147. https://doi.org/10.1093/molbev/msn059

Rohrmann, G.F., 2019. The AcMNPV genome: Gene content, conservation, and function, Baculovirus Molecular Biology [Internet]. 4th edition. National Center for Biotechnology Information (US).

Rohrmann, G.F., 2013. The baculovirus replication cycle: Effects on cells and insects. National Center for Biotechnology Information (US).

Schraiber, J.G., Evans, S.N., Slatkin, M., 2016. Bayesian Inference of Natural Selection from Allele Frequency Time Series. Genetics 203, 493–511. https://doi.org/10.1534/genetics.116.187278

Schulte, R., Makus Carsten, Hasert Barbara, Michiels Nico K., Schulenburg Hinrich, 2011. Host– parasite local adaptation after experimental coevolution of Caenorhabditis elegans and its microparasite Bacillus thuringiensis. Proc. R. Soc. B Biol. Sci. 278, 2832–2839. https://doi.org/10.1098/rspb.2011.0019

Singmann, H., Bolker, B., Westfall, J., Aust, F., Ben-Shachar, M.S., 2019. afex: Analysis of Factorial Experiments. R Package.

Vail, P.V., Tebbets, J.S., 1990. Comparative Biology and Susceptibility of Plodia interpunctella (Lepidoptera: Pyralidae) Populations to a Granulosis Virus. Environ. Entomol. 19, 791–794. https://doi.org/10.1093/ee/19.3.791

Visher, E., Boots, M., 2020. The problem of mediocre generalists: population genetics and eco-evolutionary perspectives on host breadth evolution in pathogens. Proc. R. Soc. B Biol. Sci. 287, 20201230. https://doi.org/10.1098/rspb.2020.1230

Whitlock, M.C., 1996. The Red Queen Beats the Jack-Of-All-Trades: The Limitations on the Evolution of Phenotypic Plasticity and Niche Breadth. Am. Nat. 148, S65–S77.

Wickham, H., 2009. ggplot2 – Elegant Graphics for Data Analysis. Springer.

Woolhouse, M.E.J., Gowtage-Sequeria, S., 2005. Host Range and Emerging and Reemerging Pathogens. Emerg. Infect. Dis. 11, 1842–1847. https://doi.org/10.3201/eid1112.050997

Zhan, J., Mundt, C.C., Hoffer, M.E., McDonald, B.A., 2002. Local adaptation and effect of host genotype on the rate of pathogen evolution: an experimental test in a plant pathosystem. J. Evol. Biol. 15, 634–647. https://doi.org/10.1046/j.1420-9101.2002.00428.x

