## Supplemental methods for "The Evolution of Host Specialization in an Insect Pathogen"

### Host line maintenance

Populations of these each inbred genotype were maintained in 1000mL straight-side wide-mouth Nalgene jars (ThermoFisher Scientific, U.K.) with 200 g of standard food medium in a single incubator at 27±2 °C and 35±5% humidity, with 16:8hr light:dark cycles. Standard food was made with 250g 'Ready Brek' (Weetabix Ltd., U.K.), 150g wheat bran (Bob's Red Mill, U.S.A.), 100g rice flour (Bob's Red Mill, U.S.A.), 100g brewer's yeast (MP Biomedicals, U.S.A.), 125ml glycerol (VWR, U.S.A), 125ml organic honey (Dutch Gold Honey Inc., U.S.A.), 2.2g methyl paraben (VWR, U.S.A.), and 2.2g sorbic acid (Spectrum Chemicals, U.S.A.). We mixed the food medium in batches and froze it for 24 hours before use. To maintain these populations, we moved about fifty adult moths onto fresh food jars as new adult moths emerged about monthly.

### Selection inference using genomic time series data

We used an MCMC-based inference procedure to infer the strength of selection acting at variable positions in our geomic time series data [1]. This software estimates selection coefficients given an observed frequency trajectory, accounting for uncertainty in true allele frequencies due to binomial sampling. The flexibility with respect to the number of copies of each allele detected at each locus makes this software a natural choice for our project, where the sampling is pooled across different viral particles and sequencing depth is variable across the genome. Throughout, we estimated selection strength on whichever allele was the minor allele in the ancestral viral sequences (time point 0).

We downloaded an updated version of the software from https://github.com/ekirving/selection. Our command lines used the following syntax:

```
./sr -D <input_data> -n 100000 -d 0.005 -F 20 -f 1000 -s 1000 -P <population_history> -e 8067
```

The software returns a posterior distribution of selection strength estimates and two selection coefficients, which correspond to the parameters of a full diploid selection model. We note that the virus in question here is haploid. We therefore take the average of the two selection parameters as an estimate of the selection strength [2].

Our sequence data corresponds to multiple replicates of each experiment, derived from a common ancestral stock virus sequence. This allowed us to search for repeated selection signals that are shared across replicates, as well as contingent signals that may correspond to *de novo* mutations that occurred during the experiment.

**Demographic models**

Demographic history affects patterns of genetic variation and may confound inferences of selection [3,4]. In genomic time series data, unaccounted-for changes in effective population size that occur during the experiment can result in biased estimates of selection strength[1]. This is relevant in the context of a serial infection experiment, in which individuals are inoculated with an unknown number of viral particles that then rapidly expand. While we were able to quantify the final viral load within infected individuals (which corresponds to ~$10^{10}$ particles), we did not quantify the inoculum and the precise growth kinetics within infected individuals are unknown.

Previous work has examined the population-genetics of serial passage experiments [5]. While the dynamics of serial passages can be complex, the harmonic mean of the population size during a growth epoch approximates the effective drift timescale within that timespan. We chose a range of inoculums (from 50 to 500 viral particles) and a range of growth rates (including “slow” and “fast” exponential processes with ~1.2-5 fold growth per generation) and calculated the harmonic mean of population size and the number of generations needed to reach $10^{10}$ particles for each scenario. We then repeated our estimates of selection strength using each of these effective population sizes, which ranged from small to moderate ($N_e$ = 425 to $N_e$ = 6778).

**Comparing null models of genetic drift**

We sought to understand the effects of each set of demographic assumptions by comparing the inferred selection strength under each model (i.e., each effective population size that we considered). Ideally, selection estimates for a given experimental replicate would be correlated and similar in magnitude across the demographic models, which would suggest that our results are robust to demographic model misspecification. In practice, we do not expect the estimates to be identical, because the posteriors tend to be broad suggesting substantial uncertainty in each estimate. Nonetheless, robust estimates should be correlated across demographic models.

For most variable sites, we do not expect substantial selection estimates because they are likely to be nearly neutral and drift through frequency space. Therefore, we condition on alleles that were estimated to be "significant" in the most conservative demographic model and only make comparisons to this set of alleles. We determined significance by a loose threshold, where the 90% HPD interval did not overlap 0 (see below for further justification for this threshold). We expected the most conservative demographic model to be the one with the lowest effective population size ($N_e = 425$), under which we would expect the largest random fluctuations in allele frequencies and hence the lowest (or most uncertain) selection estimates.

In Figure S3A, we compare maximum a posteriori selection strength estimates across demographic models at alleles that were called as significant in the $N_e = 425$ model. We find that selection estimates obtained under the different effective population sizes were somewhat correlated (Spearman rho=0.199, p=3.939e-06; Figure S1A), but more loci were called as significant under the higher $N_e$ models (Fig S3B). 284 of the 532 pairs of estimates in Figure S3A were significant and had the same direction of effect in both the $N_e = 425$ and the comparison model (red points in Figure S3A).

Given the variability in selection estimates across demographic models, we used the model that resulted in the fewest significant hits for downstream analyses ($N_e = 540$, which had only 114 total selection signals) to limit the potential for excessive false positive calls. Note that even with this more conservative analysis, we suggest that any selection inferences here should be taken as preliminary because we do not observe clear or repeated sweep-like signals that would be obvious indicators of strong selection. We also note that other selection processes, such as selective interference and polygenic adaptation, could underlie the frequency changes in our data and may be difficult to detect with this (or another) approach.

Estimates of selection strength for indels from across the genome are plotted in Figures S4, S5, and S6, which correspond to each of the plodia lines (2, 9, and 17, respectively). We do not plot the SNPs here.

**Repeated selection across replicates**

Among the 114 alleles that were called as significant in the analysis of the $N_e$ = 540 model, we found a small subset that were called as significant in 2 or more biological replicates with the same experimental design. In Table S5, we list each such variant. A majority of these variants were indels (13 out of 20). Only two indel variants had selection signals that were specific to a particular condition (variants at positions 27789 and 20105, both of which were detected in multiple replicates of line 17). Other putatively selected variants were shared across two or more of the Plodia lines, suggesting they may represent adaptation to the laboratory environment rather than adaptation to specific elements of each experiment. Among the 7 selection signals at SNPs, 3 were unique to specific experiments, including positions 27885, 33317, and 33358, which were detected in lines 9, 17, and 17, respectively.

In general, we note that the inferred selection coefficients are mostly indicative of weak positive selection. If we suppose an effective population size of 540, then the inferred values of *2Ns* indicate per-allele effects ranging from 0.0065 to 0.042. In Figure S7, we plot an example of observed and inferred posterior frequency trajectories for one indel that was inferred as positively selected in multiple experiments.

**Assessing evidence for sweeps and mutation accumulation**

Although we observe many instances of subtle frequency differentiation during the extent of our experiments, we observed no clear signals of selective sweeps in which a low frequency allele swept to high frequency during the experiment. Given the relatively high depth of coverage of our samples and the quality of the sequencing data, it is unlikely that we failed to detect many (if any) sweeps. However, it is possible that our experiments ended too early to capture the full timespan of some ongoing sweeps.

If complete sweeps cannot explain the specialization that was observed at the phenotype level, then more subtle frequency variation seems a likely candidate to explain the genetic mechanism for adaptation. This subtle frequency differentiation could be due to polygenic adaptation (in which frequency changes at many alleles of weak effect on a selected trait can have a substantial effect in concert) or selective interference (in which one clone fails to fix because a higher fitness clone invades and outcompetes it before fixation occurs). Either of these scenarios is not well captured by the MCMC software that we used, which is designed to detect directional selection[1].

Mutation accumulation could also play a role in specialization due to subtle frequency shifts. Under the mutation accumulation model, selection on alleles that are deleterious in another environment is relaxed in a new environment, allowing some of these alleles to drift to higher frequency and potentially alter the value of a trait that is constrained in other environments. Because mutation accumulation is a drift-based mechanism, it is not expected that the set of causal alleles will necessarily be similar across biological replicates. Without estimates of the functional effects of putative causal alleles (as might be obtained in an association study or functional assay) it is difficult to test for mutation accumulation. Given estimates of allelic effects on a trait, it might be possible to estimate the change in trait values due to frequency shifts during the experiment, which would make it possible to probe the mutation accumulation model as a potential explanation for specialization.

1. Schraiber JG, Evans SN, Slatkin M. Bayesian Inference of Natural Selection from Allele Frequency Time Series. Genetics. 2016. pp. 493–511. doi:10.1534/genetics.116.187278

2. Simonti CN, Lachance J. Ancient DNA reveals that few GWAS loci have been strongly selected during recent human history. bioRxiv. 2021. p. 2021.04.13.439742. doi:10.1101/2021.04.13.439742

3. Teshima KM. How reliable are empirical genomic scans for selective sweeps? Genome Research. 2006. pp. 702–712. doi:10.1101/gr.5105206

4. Ewing GB, Jensen JD. The consequences of not accounting for background selection in demographic inference. Mol Ecol. 2016;25: 135–141.

5. Harpak A, Sella G. Neutral null models for diversity in serial transfer evolution experiments. Evolution. 2014;68: 2727–2736.
