## Supplemental model tables for "The Evolution of Host Specialization in an Insect Pathogen"

### Model Tables for End of Evolution Proportion Infected (Figure 2a)

```
Mixed Model Anova Table (Type 3 tests, LRT-method)

Model: cbind(NumberInfected, NumberUninfected) ~ Dose + PlodiaAssayLine +
Model:     EvolvedLine + Self + (1 | EvolvedLine/VirusLine)
Data: IFDE
Df full model: 9
           Effect df      Chisq p.value
1            Dose  1 243.50 ***  <.0001
2 PlodiaAssayLine  2  16.79 ***   .0002
3     EvolvedLine  2       0.73     .69
4            Self  1     6.10 *     .01
---
Signif. codes:  0 ‘***’ 0.001 ‘**’ 0.01 ‘*’ 0.05 ‘+’ 0.1 ‘ ’ 1
```

```
Generalized linear mixed model fit by maximum likelihood (Laplace Approximation) ['glmerMod']
 Family: binomial  ( logit )
Formula: cbind(NumberInfected, NumberUninfected) ~ Dose + PlodiaAssayLine +
    EvolvedLine + Self + (1 | EvolvedLine/VirusLine)
   Data: IFDE

     AIC      BIC   logLik deviance df.resid
   287.7    305.6   -134.9    269.7       45

Scaled residuals:
    Min      1Q  Median      3Q     Max
-2.2570 -0.8435  0.2470  0.6466  4.3307

Random effects:
 Groups                Name        Variance  Std.Dev.
 VirusLine:EvolvedLine (Intercept) 5.107e-02 2.260e-01
 EvolvedLine           (Intercept) 3.002e-10 1.733e-05
Number of obs: 54, groups:  VirusLine:EvolvedLine, 9; EvolvedLine, 3

Fixed effects:
                   Estimate Std. Error z value Pr(>|z|)
(Intercept)       -2.343429   0.238096  -9.842  < 2e-16 ***
Dose               0.045898   0.003276  14.010  < 2e-16 ***
PlodiaAssayLine9  -0.213720   0.170298  -1.255   0.2095
PlodiaAssayLine17 -0.635957   0.163102  -3.899 9.65e-05 ***
EvolvedLine9       0.208154   0.247733   0.840   0.4008
EvolvedLine17      0.061621   0.247485   0.249   0.8034
SelfTRUE           0.339872   0.137486   2.472   0.0134 *
---
Signif. codes:  0 ‘***’ 0.001 ‘**’ 0.01 ‘*’ 0.05 ‘.’ 0.1 ‘ ’ 1
```

### Model Tables for End of Evolution Ave Number of Virions Produced (Figure 2b)

```
Mixed Model Anova Table (Type 3 tests, LRT-method)

Model: AveCount ~ PlodiaAssayLine + EvolvedLine + Self + (1 | EvolvedLine/VirusLine)
Data: IFDEMD
Df full model: 8
           Effect df      Chisq p.value
1 PlodiaAssayLine  2 3141.10 ***  <.0001
2     EvolvedLine  2    7.02 *      .03
3            Self  1 3776.50 ***  <.0001
---
Signif. codes:  0 ‘***’ 0.001 ‘**’ 0.01 ‘*’ 0.05 ‘+’ 0.1 ‘ ’ 1
```

```
Generalized linear mixed model fit by maximum likelihood (Laplace Approximation) ['glmerMod']
 Family: poisson  ( log )
Formula: AveCount ~ PlodiaAssayLine + EvolvedLine + Self + (1 | EvolvedLine/VirusLine)
   Data: IFDEMD

     AIC      BIC   logLik deviance df.resid
 10493.3  10503.6  -5238.6  10477.3       19

Scaled residuals:
    Min      1Q  Median      3Q     Max
-28.298 -14.961  -6.147  12.157  51.546

Random effects:
 Groups                Name        Variance  Std.Dev.
 VirusLine:EvolvedLine (Intercept) 7.504e-02 2.739e-01
 EvolvedLine           (Intercept) 2.246e-10 1.499e-05
Number of obs: 27, groups:  VirusLine:EvolvedLine, 9; EvolvedLine, 3

Fixed effects:
                  Estimate Std. Error z value Pr(>|z|)
(Intercept)        6.37270    0.15876  40.141   <2e-16 ***
PlodiaAssayLine9   0.34308    0.01532  22.390   <2e-16 ***
PlodiaAssayLine17  0.74867    0.01390  53.876   <2e-16 ***
EvolvedLine9      -0.33686    0.22425  -1.502   0.1331
EvolvedLine17      0.49166    0.22407   2.194   0.0282 *
SelfTRUE           0.68261    0.01098  62.165   <2e-16 ***
---
Signif. codes:  0 ‘***’ 0.001 ‘**’ 0.01 ‘*’ 0.05 ‘.’ 0.1 ‘ ’ 1
```

**Model Tables for End of Evolution Fitness (Figure 2c)**

```
Mixed Model Anova Table (Type 3 tests, LRT-method)

Model: VirFit ~ PlodiaAssayLine + EvolvedLine + Self + (1 | EvolvedLine/VirusLine)
Data: IFDEMD
Df full model: 8
           Effect df      Chisq p.value
1 PlodiaAssayLine  2   78.75 ***  <.0001
2     EvolvedLine  2       4.32     .12
3            Self  1 1628.28 ***  <.0001
---
Signif. codes:  0 ‘***’ 0.001 ‘**’ 0.01 ‘*’ 0.05 ‘+’ 0.1 ‘ ’ 1
```

```
Generalized linear mixed model fit by maximum likelihood (Laplace Approximation) ['glmerMod']
 Family: poisson  ( log )
Formula: VirFit ~ PlodiaAssayLine + EvolvedLine + Self + (1 | EvolvedLine/VirusLine)
   Data: IFDEMD

     AIC      BIC   logLik deviance df.resid
  4913.4   4923.8  -2448.7   4897.4       19

Scaled residuals:
     Min       1Q   Median       3Q      Max
-18.9832  -9.6798  -0.1794   7.2976  27.4487

Random effects:
 Groups                Name        Variance  Std.Dev.
 VirusLine:EvolvedLine (Intercept) 8.016e-02 2.831e-01
 EvolvedLine           (Intercept) 5.838e-10 2.416e-05
Number of obs: 27, groups:  VirusLine:EvolvedLine, 9; EvolvedLine, 3

Fixed effects:
                  Estimate Std. Error z value Pr(>|z|)
(Intercept)        5.97536    0.16453  36.318  < 2e-16 ***
PlodiaAssayLine9   0.08020    0.02070   3.875 0.000107 ***
PlodiaAssayLine17  0.17474    0.01977   8.840  < 2e-16 ***
EvolvedLine9      -0.28690    0.23225  -1.235 0.216713
EvolvedLine17      0.26383    0.23200   1.137 0.255458
SelfTRUE           0.65310    0.01603  40.753  < 2e-16 ***
---
Signif. codes:  0 ‘***’ 0.001 ‘**’ 0.01 ‘*’ 0.05 ‘.’ 0.1 ‘ ’ 1
```

**Model Table for End of Evolution Interaction between Virus Line and Self**

```
Mixed Model Anova Table (Type 3 tests, LRT-method)

Model: VirFit ~ PlodiaAssayLine + VirusLine * Self + (1 | EvolvedLine/VirusLine)
Data: IFDEMD
Df full model: 22
           Effect df     Chisq p.value
1 PlodiaAssayLine  2  233.22 ***  <.0001
2       VirusLine  8   56.84 ***  <.0001
3            Self  1 1122.01 ***  <.0001
4  VirusLine:Self  8 1080.59 ***  <.0001
---
Signif. codes:  0 ‘***’ 0.001 ‘**’ 0.01 ‘*’ 0.05 ‘+’ 0.1 ‘ ’ 1
```

**Model Table for Specialization over Time Series (Figure 3)**

```
Mixed Model Anova Table (Type 3 tests, LRT-method)

Model: VirFit ~ PlodiaAssayLine + Self + EvolvedLine + PassageNumber +
Model:     Self:PassageNumber + (1 | EvolvedLine/VirusLine/PassageNumber)
Data: IFDMD
Df full model: 17
              Effect df     Chisq p.value
1    PlodiaAssayLine  2 6084.68 ***  <.0001
2               Self  1  199.15 ***  <.0001
3        EvolvedLine  2       4.02     .13
4      PassageNumber  4   28.89 ***  <.0001
5 Self:PassageNumber  4 3032.20 ***  <.0001
---
Signif. codes:  0 ‘***’ 0.001 ‘**’ 0.01 ‘*’ 0.05 ‘+’ 0.1 ‘ ’ 1
[illegible]
```

```
Generalized linear mixed model fit by maximum likelihood (Laplace Approximation) ['glmerMod']
 Family: poisson  ( log )
Formula: VirFit ~ PlodiaAssayLine + EvolvedLine + Self * PassageNumber +
    (1 | EvolvedLine/VirusLine/PassageNumber)
   Data: IFDMD

     AIC      BIC   logLik deviance df.resid
 40028.4  40076.9 -19997.2  39994.4      111
```

```
Scaled residuals:
    Min      1Q  Median      3Q     Max
-38.198  -9.731  -4.146  10.068  52.631

Random effects:
 Groups                             Name        Variance  Std.Dev.
 PassageNumber:(VirusLine:EvolvedLine) (Intercept) 4.934e-01 7.024e-01
 VirusLine:EvolvedLine              (Intercept) 8.266e-10 2.875e-05
 EvolvedLine                        (Intercept) 4.007e-10 2.002e-05
Number of obs: 128, groups:
PassageNumber:(VirusLine:EvolvedLine), 45; VirusLine:EvolvedLine, 9; EvolvedLine, 3

Fixed effects:
                          Estimate Std. Error z value Pr(>|z|)
(Intercept)              7.367e+00  2.772e-01  26.578  < 2e-16 ***
PlodiaAssayLine9         2.346e-02  6.877e-03   3.411 0.000646 ***
PlodiaAssayLine17       -5.353e-01  8.119e-03 -65.934  < 2e-16 ***
EvolvedLine9            -2.255e-01  2.568e-01  -0.878 0.379842
EvolvedLine17            3.096e-01  2.569e-01   1.205 0.228160
SelfTRUE                 3.545e-05  1.100e-02   0.003 0.997428
PassageNumber1          -1.957e+00  3.320e-01  -5.894 3.76e-09 ***
PassageNumber4          -6.022e-01  3.313e-01  -1.818 0.069071 .
PassageNumber6          -6.178e-01  3.312e-01  -1.865 0.062155 .
PassageNumber9          -1.263e+00  3.313e-01  -3.812 0.000138 ***
SelfTRUE:PassageNumber1 -5.818e-02  2.585e-02  -2.250 0.024424 *
SelfTRUE:PassageNumber4 -3.811e-01  1.861e-02 -20.473  < 2e-16 ***
SelfTRUE:PassageNumber6  1.823e-01  1.725e-02  10.571  < 2e-16 ***
SelfTRUE:PassageNumber9  7.805e-01  1.953e-02  39.965  < 2e-16 ***
---
Signif. codes:  0 ‘***’ 0.001 ‘**’ 0.01 ‘*’ 0.05 ‘.’ 0.1 ‘ ’ 1
```

**Model Table for Correlations between Viral Productivity and Infectivity (Figure 4)**

```
Mixed Model Anova Table (Type 3 tests, LRT-method)

Model: AveCount ~ PlodiaAssayLine + EvolvedLine + Self * PropInf * PassageNumber +
Model:     (1 | EvolvedLine/VirusLine/PassageNumber)
Data: IFDMD
Df full model: 27
                      Effect df       Chisq p.value
1            PlodiaAssayLine  2  864.80 ***  <.0001
2                EvolvedLine  2      7.46 *     .02
3                       Self  1  141.97 ***  <.0001
4                    PropInf  1 2523.48 ***  <.0001
5              PassageNumber  4   42.60 ***  <.0001
6               Self:PropInf  1   41.91 ***  <.0001
7         Self:PassageNumber  4 9222.52 ***  <.0001
8      PropInf:PassageNumber  4 3948.49 ***  <.0001
9 Self:PropInf:PassageNumber  4 6716.85 ***  <.0001
---
Signif. codes:  0 ‘***’ 0.001 ‘**’ 0.01 ‘*’ 0.05 ‘+’ 0.1 ‘ ’ 1
```

```
Generalized linear mixed model fit by maximum likelihood (Laplace Approximation) ['glmerMod']
 Family: poisson  ( log )
Formula: AveCount ~ PlodiaAssayLine + EvolvedLine + Self * PropInf * PassageNumber +
    (1 | EvolvedLine/VirusLine/PassageNumber)
   Data: IFDMD

     AIC      BIC   logLik deviance df.resid
 38695.3  38771.2 -19320.6  38641.3       96

Scaled residuals:
    Min      1Q  Median      3Q     Max
-58.107  -7.829   0.082   7.444  60.577

Random effects:
 Groups                                Name        Variance  Std.Dev.
 PassageNumber:(VirusLine:EvolvedLine) (Intercept) 2.643e-01 5.141e-01
 VirusLine:EvolvedLine                 (Intercept) 2.826e-10 1.681e-05
 EvolvedLine                           (Intercept) 2.438e-10 1.561e-05
Number of obs: 123, groups:
PassageNumber:(VirusLine:EvolvedLine), 45; VirusLine:EvolvedLine, 9; EvolvedLine, 3
```

```
Fixed effects:
                                Estimate Std. Error z value Pr(>|z|)    
(Intercept)                     7.295994   0.204046  35.757  < 2e-16 ***
PlodiaAssayLine9                0.077622   0.005029  15.433  < 2e-16 ***
PlodiaAssayLine17               0.206107   0.007032  29.309  < 2e-16 ***
EvolvedLine9                   -0.206584   0.187937  -1.099 0.271673    
EvolvedLine17                   0.472538   0.187917   2.515 0.011916 *  
SelfTRUE                       -0.001390   0.039242  -0.035 0.971746    
PropInf                         0.498062   0.034479  14.445  < 2e-16 ***
PassageNumber1                 -0.401076   0.244680  -1.639 0.101174    
PassageNumber4                  0.484948   0.243355   1.993 0.046288 *  
PassageNumber6                 -0.818059   0.244320  -3.348 0.000813 ***
PassageNumber9                 -1.395389   0.245410  -5.686  1.3e-08 ***
SelfTRUE:PropInf                0.002351   0.064585   0.036 0.970958    
SelfTRUE:PassageNumber1        -1.550782   0.061801 -25.093  < 2e-16 ***
SelfTRUE:PassageNumber4        -1.109339   0.043075 -25.754  < 2e-16 ***
SelfTRUE:PassageNumber6         0.296769   0.048279   6.147  7.9e-10 ***
SelfTRUE:PassageNumber9         3.445675   0.067575  50.991  < 2e-16 ***
PropInf:PassageNumber1         -1.487333   0.064583 -23.030  < 2e-16 ***
PropInf:PassageNumber4          0.110412   0.041350   2.670 0.007581 ** 
PropInf:PassageNumber6          3.799742   0.079924  47.542  < 2e-16 ***
PropInf:PassageNumber9          1.138958   0.076284  14.931  < 2e-16 ***
SelfTRUE:PropInf:PassageNumber1  3.010896   0.121183  24.846  < 2e-16 ***
SelfTRUE:PropInf:PassageNumber4  2.772271   0.081694  33.935  < 2e-16 ***
SelfTRUE:PropInf:PassageNumber6 -1.415432   0.109270 -12.954  < 2e-16 ***
SelfTRUE:PropInf:PassageNumber9 -5.647893   0.128738 -43.871  < 2e-16 ***
---
Signif. codes:  0 ‘***’ 0.001 ‘**’ 0.01 ‘*’ 0.05 ‘.’ 0.1 ‘ ’ 1
```
