## Supplemental Figure for "The Evolution of Host Specialization in an Insect Pathogen"

### Supplemental Tables and Figures

#### Supplemental Figure 1: Passage and Assay Scheme

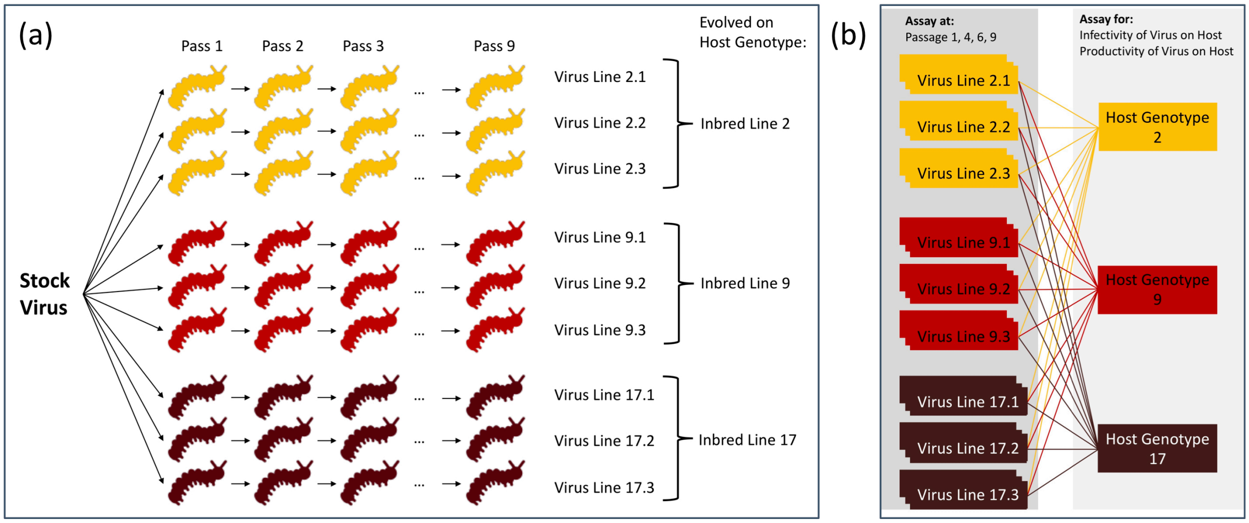


**Figure S1.** (a) Passaging scheme for virus experimental evolution lines. Stock virus was used to initiate nine virus evolution lines, 3 per host genotype. Virus was serially passaged through larvae for 9 passages. (b) Assay scheme for virus experimental evolution lines. The starting virus stock and every virus line at passages 1, 4, 6, and 9 was assayed for infectivity and productivity on each host genotype.

#### Supplemental Table 1: Resistance of Host Genotypes

| | Line 2 | Line 17 | Line 9 |
|---|---|---|---|
| Proportion infected at 5.6 x 10^8 OB/mL Dilution | 15/19 | 7/11 | 7/9 |
| Percent | 78.9% | 63.6% | 77.8% |
| Proportion infected at 2.8 x10^8 OB/mL Dilution | 5/18 | 6/15 | 10/20 |
| Percent | 27.8% | 40% | 50% |

The passage dose was ~7.5x10^8 occlusion bodies per mL.

**Supplemental Table 2: Passaging Conditions for Cadavers Harvested and Virus Counts**

| # Infected Individuals Harvested | | | | | | | | | |
|---|---|---|---|---|---|---|---|---|---|
| Virus Line | P1 | P2 | P3 | P4 | P5 | P6 | P7 | P8 | P9 |
| 2.1 | 10 | 10 | 10 | 10 | 9 | 10 | 10 | 10 | 10 |
| 2.2 | 10 | 10 | 5 | 10 | 10 | 10 | 10 | 10 | 10 |
| 2.3 | 10 | 8 | 2 | 10 | 10 | 5 | 10 | 10 | 10 |
| 9.1 | 10 | 10 | 9 | 10 | 10 | 5 | 10 | 10 | 10 |
| 9.2 | 10 | 10 | 4 | 10 | 10 | 8 | 10 | 10 | 10 |
| 9.3 | 10 | 10 | 10 | 10 | 10 | 5 | 10 | 10 | 10 |
| 17.1 | 10 | 10 | 7 | 10 | 10 | 10 | 10 | 10 | 10 |
| 17.2 | 10 | 10 | 10 | 10 | 10 | 10 | 10 | 10 | 10 |
| 17.3 | 10 | 10 | 4 | 10 | 8 | 9 | 10 | 10 | 10 |

| Virus Count | | | | | | | | | |
|---|---|---|---|---|---|---|---|---|---|
| Virus Line | P1 | P2 | P3 | P4 | P5 | P6 | P7 | P8 | P9 |
| 2.1 | 265 | | | 5200 | 7450 | 6700 | 11000 | 4850 | 4650 |
| 2.2 | 3200 | | | 6825 | 8400 | 6025 | 4775 | 7700 | 6363 |
| 2.3 | | | | 3975 | 3000 | 1850 | 8650 | 5725 | 4800 |
| 9.1 | 6250 | | | 2950 | 3300 | 3700 | 5350 | 4850 | 5025 |
| 9.2 | 2525 | | | 9425 | 3925 | 2525 | 3700 | 3300 | 2200 |
| 9.3 | | | | 5700 | 3575 | 2175 | 6375 | 5850 | 4475 |
| 17.1 | 6475 | | | 3550 | 3175 | 1275 | 1E+05 | 3225 | 5988 |
| 17.2 | 2700 | | | 2900 | 3900 | 8200 | 4950 | 7250 | 4425 |
| 17.3 | | | | 7525 | 2000 | 6950 | 8200 | 4025 | 4863 |

Counting was done by diluting the solution to a countable dose, adding 5uL to the Petroff Hauser counting chamber and counting virus under darkfield microscopy at 400x. Four .02 cubic mm squares were counted, then they were averaged and multiplied by the dilution factor for the counts here.

Particles per mL = count*25/.02*1000

Passaging concentration was 600 count

#### Supplemental Table 3: Methods and Results for Virus Purification Method Test

**Introduction:** We wanted to ensure that we were not passaging virus alongside other microbes that might contaminate our experiment. While we did not observe mortality from other infections at any point during out experimental evolution, we devised a novel method to semi-purify baculovirus.

**Methods:** We collected 10 healthy larvae from populations un-exposed to PiGV and 10 larvae that had been killed by PiGV. We extracted virus from these pooled cadavers by tissue homogenization then transferred 1mL of the supernatant to a sterile 1.5mL Eppendorf tube and centrifuged the solution for 1 minute at 3,000 rpm to remove larger particulate matter from the supernatant. We transferred 600uL of this solution to a sterile 1.5 mL Eppendorf and centrifuged this for 3 minutes at 13,000 rpm to pellet the virus. We removed the supernatant from the pellet and resuspended in 1mL sterile water. We then diluted this solution 10x. For treatments with a filtration step, we added 600uL of the 10x solution to .65 or .45 micron filter spin columns in triplicate (Millipore Sigma, U.S.A.) that we centrifuged at 13,000rpm for 3 minutes. For unfiltered (2), .65 micron filtered (3), or .45 micron filtered (3) samples, we then plated 1mL of the solution onto both King's Medium B agar plates (bacteria) and Sabouraud agar plates (fungus) and incubated plates for 5 days at 27C. Growth on the plates was then recorded. Virus titers from each treatment were also counted on a Petroff Hauser counting chamber (as elsewhere) to determine yield. Based on it's acceptable yields and moderate bacteria and fungal filtering, we chose to continue with the .65 micron filter.

**Results**

| | | Kings Medium B agar plate | Sabouraud agar plate | Virus Yield |
|---|---|---|---|---|
| Homogenate from healthy larvae | Not Filtered | lawn | many spots | 0 |
| | | lawn | many spots | |
| | .65 micron filtered | no growth | no growth | 0 |
| | | no growth | no growth | |
| | | no growth | no growth | |
| | .45 micron filtered | 1 spot | no growth | 0 |
| | | no growth | no growth | |
| | | no growth | no growth | |
| Homogenate from PiGV infected larvae | Not Filtered | lawn | 4 colonies | 100% |
| | | lawn | 4 colonies | |
| | .65 micron filtered | lawn | no growth | 41.25% |
| | | small red and white plaques | 1 colony | |
| | | 2 dots | 1 colony | |
| | .45 micron filtered | no growth | 1 colony | 12.46% |
| | | no growth | no growth | |
| | | no growth | no growth | |

#### Supplemental Figure 2 – Evolution of Specialization across Time Series

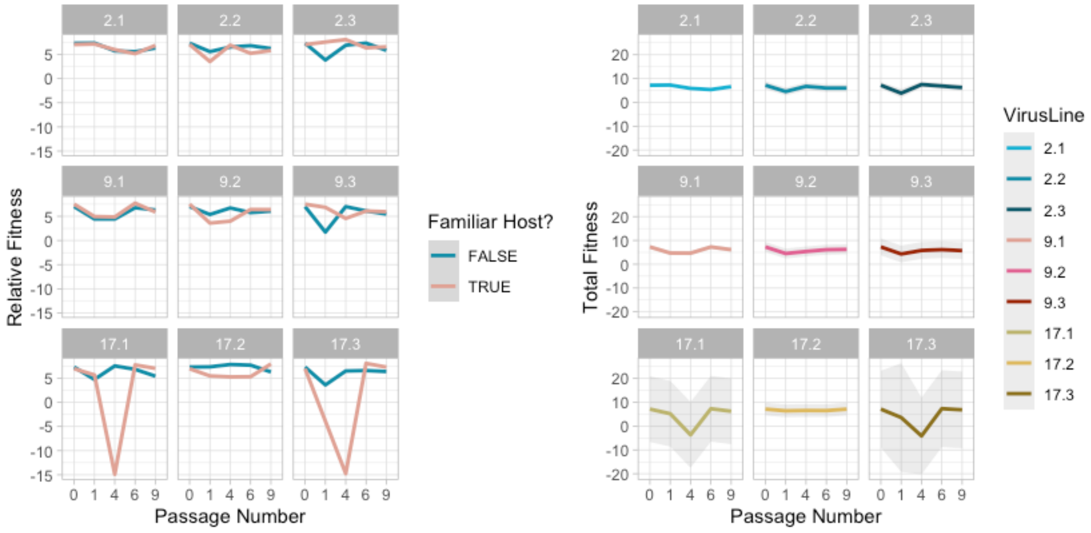


**Figure S2**-Evolution of Specialization over Time. Panelled plot showing the effect of whether the virus was assayed on its familiar host genotype (red) or on a foreign one (yellow) on viral (a) infectivity, (b) productivity, and (c) fitness over time. Virus lines evolve significantly higher relative infectivity, productivity, and fitness on familiar lines over the experiment. Effects are taken from the GLMM models using the 'emmeans' package.

#### Supplemental Table 4: Function Annotation for Regions of High Variation

| Position(s) | Gene and homologs | Known function of homolog |
|---|---|---|
| 12986 | PiGV ORF15; similar to AcMNPV ac11 | needed for budded and occluded production |
| 17001 | pep-1; PiGV ORF19; similar to CpGV cp20 and AcMNPV ac131 | OB stability |
| 17266-18168 | PiGV ORF20; similar to CrleGV-CV3 ORF21 and AcMNPV ac18 | longer time to kill ; involved in oral infection |
| 23228-23704 | PiGV ORF25 | |
| 26418 | efp; PiGV ORF27; similar to CpGV cp31 and AcMNPV ac23 | cell binding/infection/time to kill/productivity |
| 27744-28643 | PiGV ORF28 | |
| 33269-33358 | odv-e66; PiGV ORF32; similar to CpGV cp37 and AcMNPV ac46 | may be involved in the digestion of the peritrophic matrix |
| 34193 | lef-2; PiGV ORF34; similar to CpGV cp41 and AcMNPV ac6 | DNA replication |
| 37127 | mmp; PiGV ORF38; similar to CpGV cp46 | may be involved in the digestion of the peritrophic matrix |
| 42427 | PiGV ORF43; similar to CpGV cp50 | |
| 49848 | PiGV ORF53 | |
| 51501-51544 | PiGV ORF54 | |

| | | |
|---|---|---|
| 54460 | p24; PiGV ORF58; similar to CpGV cp71 and AcMNPV ac129 | longer time to kill |
| 56299 | pif-1; PiGV ORF62; similar to CpGV cp75 and AcMNPV ac119 | needed for oral infection |
| 58015 | fgf-1; PiGV ORF64; similar to CpGV cp76 and AcMNPV ac32 | entering nucleus |
| 61600 | p45; p48; PiGV ORF70; similar to CpGV cp83 and AcMNPV ac103 | virus release from cell |
| 65827-65830 | 38k; PiGV ORF75; similar to CpGV cp88 and AcMNPV ac98 | nucleocapsid formation |
| 67548 | PiGV ORF77; similar to CpGV cp90 and AcMNPV ac95 | implicated in host range |
| 69992-70710 | odv-e25; PiGV ORF78; similar to CpGV cp91 and AcMNPV ac94 | virus occlusion |
| 74133 | lef-4; PiGV ORF82; similar to CpGV cp95 and AcMNPV ac90 | translation? |
| 99853 | PiGV ORF111; similar to CpGV cp12 | |
| 103038 | rr2a; PiGV ORF113; similar to CpGV cp128 | ribonucleotide reductase |
| 106555 | na | |
| 106601-106604 | na | |
| 108237 | PiGV ORF118; similar to CpGV cp135 | |
| | AcMNPV homolog functions are from Rohrmann (2019). https://www.ncbi.nlm.nih.gov/books/NBK543457/ | |

#### Supplemental Figure 3

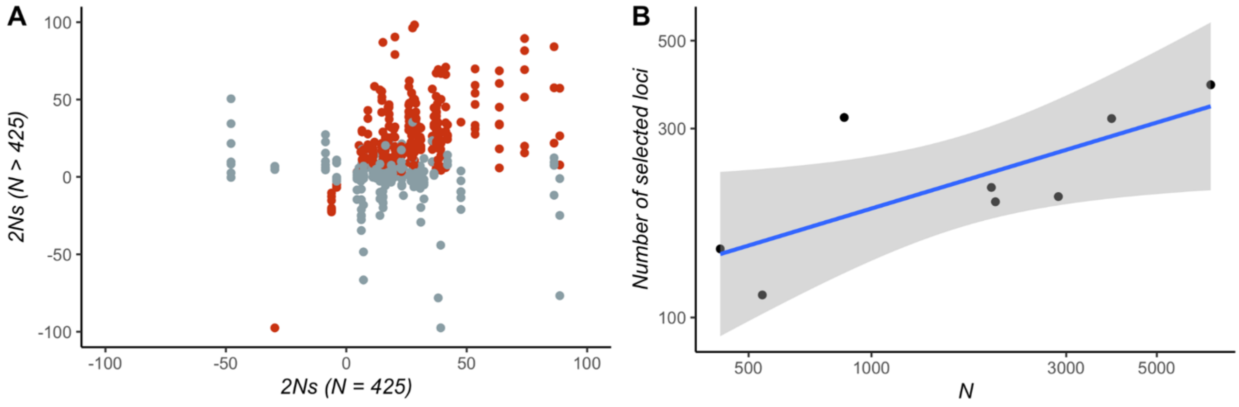


**Figure S3: A.** Relationship between estimates of selection strength obtained under different drift models, with the *N*=425 model on the x-axis and all other models plotted on the y-axis. We observed a modest but significant correlation (Spearman rho=0.199, p=3e-6) between the estimates. Note that some large selection strength estimates fell outside this range and are not plotted. The red points had significant selection estimates in both models. Loci were considered significant if the 90% HPD region of the selection coefficient posterior excluded 0. **B.** The number of significantly selected loci in each drift model. The blue line represents a simple linear model fit to the points with the function `geom_smooth(method="lm")` in `ggplot2`.

#### Supplemental Figure 4

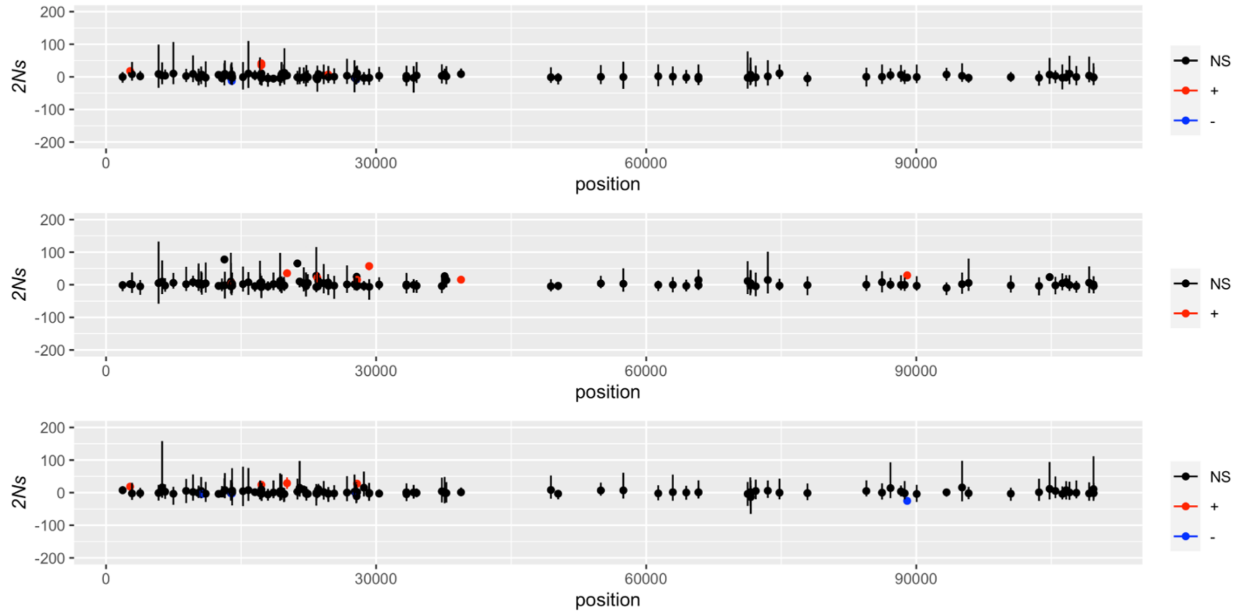


**Figure S4**: Genome-wide selection inferences of indel variants for each of the three replicates of line 2. Significantly positive fitness-increasing variants are in red, negative in blue, and non-significant in black.

#### Supplemental Figure 5

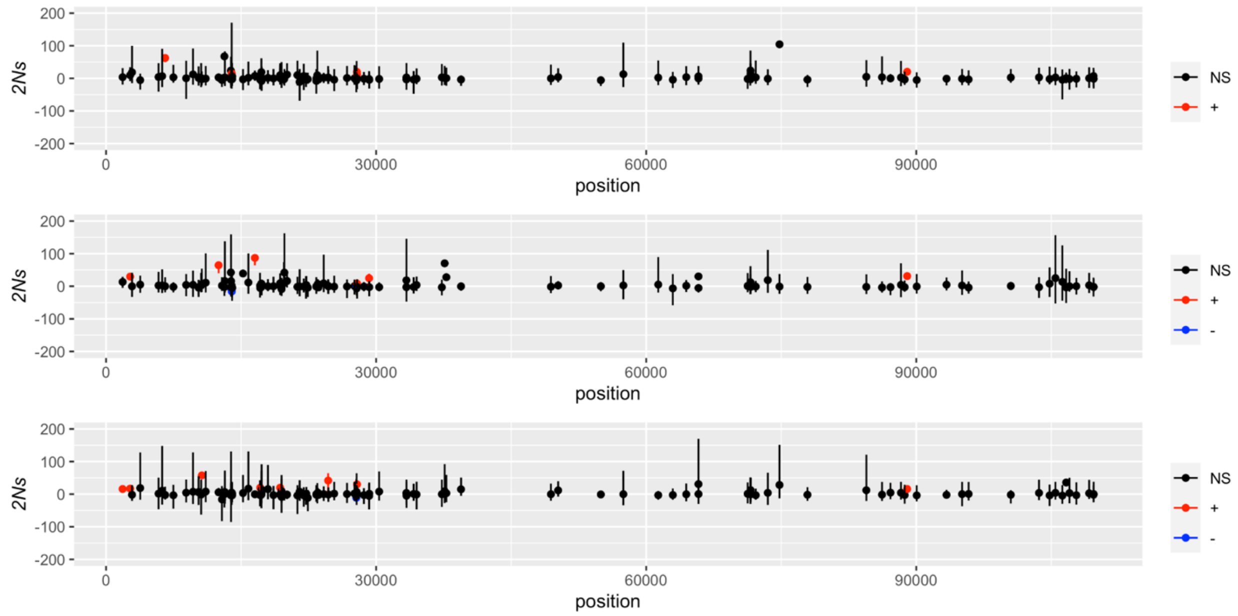


**Figure S5**: Genome-wide selection inferences of indel variants for each of the three replicates of line 9. Significantly positive fitness-increasing variants are in red, negative in blue, and non-significant in black.

#### Supplemental Figure 6

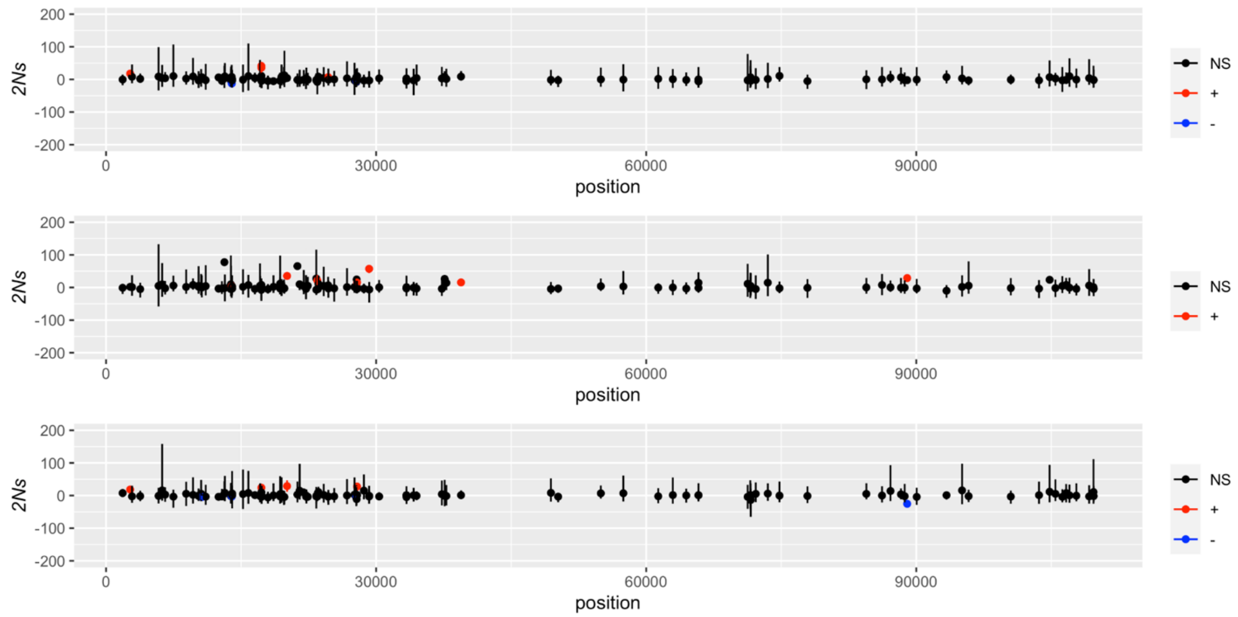


**Figure S6**: Genome-wide selection inferences of indel variants for each of the three replicates of line 17. Significantly positive fitness-increasing variants are in red, negative in blue, and non-significant in black.

#### Supplemental Figure 7

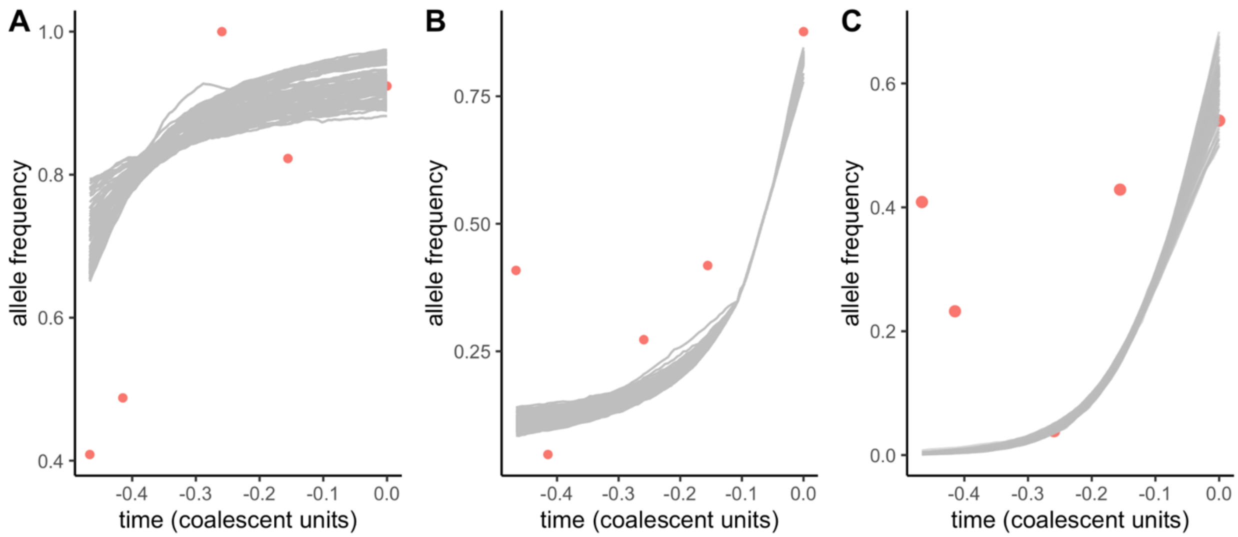


**Figure S7:** Posterior fits (gray lines) and observed frequencies (red) for indel 88992 in experimental line 9. Note that the depth of sequencing is variable and hence some of the frequency estimates are highly uncertain

#### Supplemental Table 5

| Position | Type | *2Ns* estimates | Experiment | Replicates |
|---|---|---|---|---|
| 13964 | INDEL | -13.8 -2.8 | 2 | 2 |
| 13964 | INDEL | -12.6 -3.2 | 17 | 2 |
| 20105 | INDEL | 35.4 29.0 | 17 | 2 |
| 2674 | INDEL | 29.5 17.0 | 9 | 2 |
| 2674 | INDEL | 17.8 18.5 | 17 | 2 |
| 27875 | INDEL | 13.6 45.6 | 2 | 2 |
| 27875 | INDEL | 12.4 8.4 30.2 | 9 | 3 |
| 27875 | INDEL | 15.3 27.7 | 17 | 2 |
| 27813 | INDEL | 7.8 14.6 | 2 | 2 |
| 27813 | INDEL | -5.8 -2.7 | 17 | 2 |
| 88992 | INDEL | 19.7 24.0 | 2 | 2 |
| 88992 | INDEL | 20.1 31.1 15.9 | 9 | 3 |
| 27789 | INDEL | -5.5 -4.7 | 17 | 2 |
| 108237 | SNP | 19.3 7.6 44.0 | 2 | 3 |
| 108237 | SNP | 20.6 32.0 | 9 | 2 |
| 33316 | SNP | 9.6 15.2 16.2 | 2 | 3 |
| 33316 | SNP | 18.6 16.9 15.0 | 17 | 3 |
| 27885 | SNP | 12.2 7.0 | 9 | 2 |
| 33317 | SNP | 15.9 8.8 | 17 | 3 |
| 33358 | SNP | 9.4 20.0 | 17 | 2 |

**Table S5**: Variants with consistent selection signals in 2 or more biological replicates (*i.e.*, at least 2 selection inferences where the 90% HPD did not overlap 0). The inferred values of *2Ns* are given in the middle column, and the number of replicates with significant selection signals is given in the right hand column.

#### Supplemental Figure 8: SNP and Indel frequencies across the genome

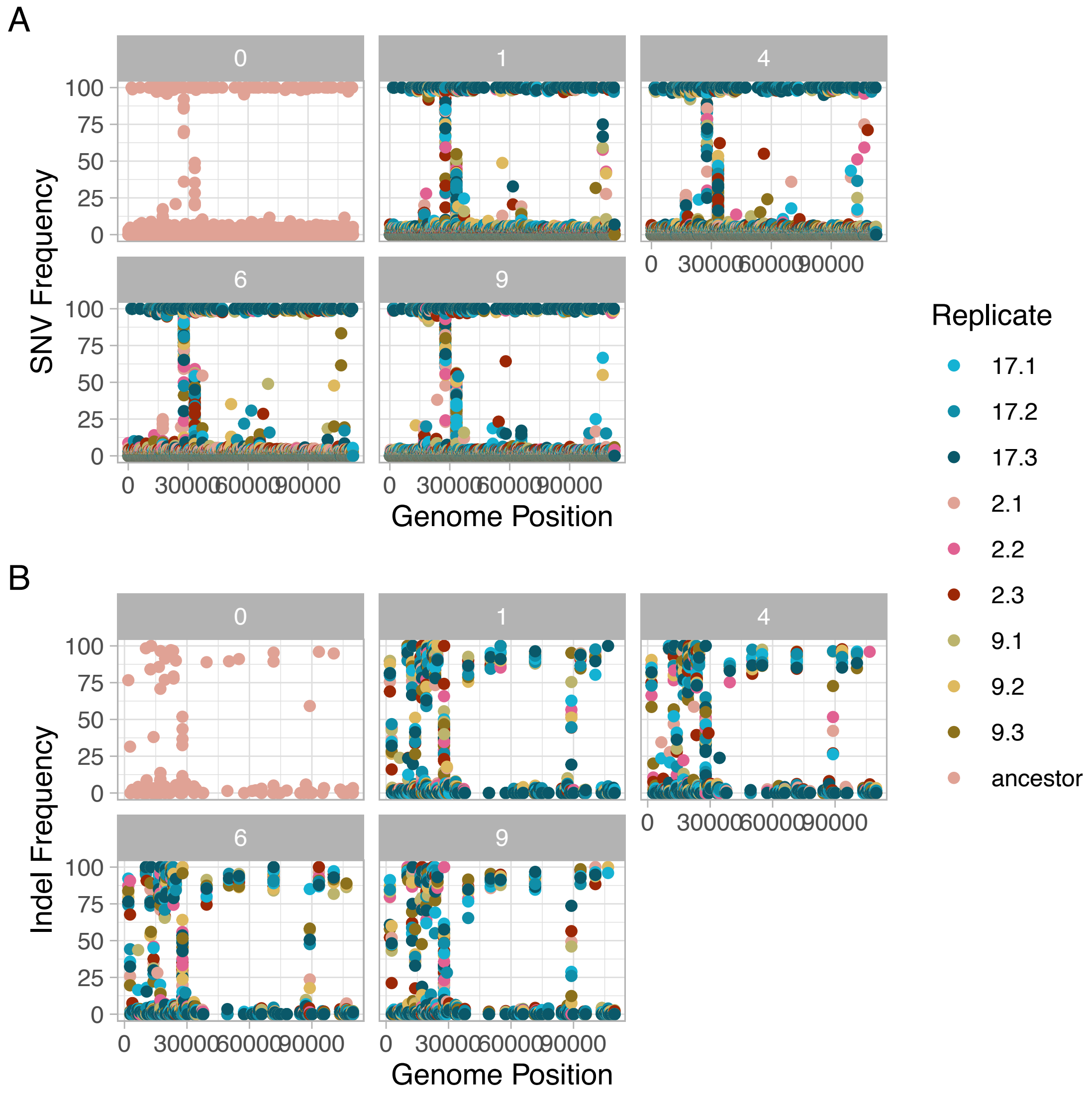


**Figure S8: A.** SNP and **B.** Indel frequencies across the genome.
